# Peptide-RNA photocrosslinks with tunable RNA chain map RNA interfaces proteome-wide

**DOI:** 10.1101/2025.09.05.674461

**Authors:** Shuyao Sha, Bernhard Kuster, Jakob Trendel

## Abstract

Photocrosslinking mass spectrometry (MS) can map protein-RNA interfaces in living cells, but current isolation procedures for peptide-RNA crosslinks eliminate most RNA and therefore cannot identify the crosslinked RNA sequence. We introduce pepR-MS, a method that enriches peptide-RNA crosslinks with RNA chains of tunable length. To maximize proteomic depth, we applied pepR-MS to generate highly truncated RNA adducts, identifying over 34,000 unique crosslinks in 589 proteins of breast cancer cells and pinpointing RNA contacts in hundreds of structured domains and intrinsically disordered regions. By modulating nuclease digestion, longer RNA chains of up to five nucleotides were retained and sequenced alongside the crosslinked peptide using a specialized MS strategy. For RNA sequence assignment, we further introduce the computational tool SALT, which identified peptide-RNA sequence pairs for over 1,600 crosslinks in a single LC-MS run. Together, pepR-SALT provides a framework for mapping protein-RNA interfaces across the proteome and transcriptome at single-amino acid and nucleotide resolution.

## Introduction

In the human proteome, over 3,000 proteins have been reported to interact with RNA^1^. About 60% of these contain one or more canonical RNA-binding domains, whereas many of the remainder interact with RNA via intrinsically disordered regions (IDRs). Elucidating these contacts at molecular resolution is crucial for understanding their biological functions^2^.

Photocrosslinking can be used to fix protein and RNA interactions in living cells, where UV irradiation covalently crosslinks proteins in direct contact with RNA. Crosslinking efficiency is typically low but can be boosted by metabolic labelling of the transcriptome with photoactivatable nucleotides such as 4-thiouridine (4SU)^3^. Photocrosslinked protein-RNA complexes can be specifically enriched for analysis of either their RNA or protein components. Crosslinking and immunoprecipitation followed by sequencing (CLIP-seq^3–6^) isolates individual proteins and uses RNA sequencing to map their interactions across the transcriptome. Conversely, RNA-contacting peptides can be co-purified with mRNA or total RNA and analysed by liquid chromatography-mass spectrometry (LC-MS), mapping RNA interaction sites across the proteome^7–11^. However, purification of photocrosslinked peptide-RNA crosslinks remains difficult due to their low abundance and bipartite composition. Crosslinking can occur at any amino acid, creating numerous combinations of modification sites and RNA adduct masses that complicate peptide identification by conventional search software. Early strategies therefore circumvented direct analysis of the crosslinked peptide but inferred RNA-interacting regions from neighbouring non-crosslinked peptides^12,13^, or from peptides depleted after irradiation^14,15^. First approaches that purified the actual peptide-RNA crosslinks used endpoint nuclease digestion to leave only up to three nucleotides attached to the peptides, thereby reducing complexity and facilitating MS search^7–11^. Studies that directly detected RNA adducts reported no more than 400 unique peptides from roughly 100 human proteins. Notably, these RNA mass offsets did not include sequence information but only the composition of the crosslinked RNA moiety. These modest yields led to the assumption that the large number of possible peptide-RNA combinations, even with short RNA chains, exceeded the capacity of MS search software at the time, making it impossible to detect more than the most abundant peptide-RNA crosslinks^16^. A pragmatic solution was the complete removal of the RNA chain from peptide-RNA crosslinks by chemical digestion with hydrofluoric acid (HF), which reliably cleaves RNA to a single nucleotide or only the crosslinked base. This approach mapped approximately 2,000 RNA binding sites in 600 human proteins using 254-nm UV irradiation^16^, and 1,500 RNA binding sites in around 300 proteins using metabolic labelling with 4SU and 365-nm UV irradiation^17^. While this represented a sizable improvement over previous approaches, removing the entire RNA chain made sequencing both the peptide and RNA components of photocrosslinked protein-RNA interfaces categorically impossible.

Here, we identify residual non-crosslinked RNA as a major cause of poor LC-MS detection of peptide-RNA crosslinks in nuclease-based workflows. We present an improved workflow termed pepR-MS (peptide-RNA crosslink isolation for sequencing by MS) that removes non-crosslinked RNA to a similar degree as chemical digestion with HF while preserving an RNA chain of adjustable length on the crosslinked peptide.

We employ pepR-MS in combination with nuclease P1 digestion to generate short-chain peptide-RNA crosslinks, which are easily detected and map thousands of RNA interaction sites across the proteome of breast cancer cells in a single MS run. With benzonase digestion, we retain longer crosslinked RNA chains that can be sequenced together with the peptide using a dual-fragmentation MS strategy. To analyse these data, we introduce the search tool SALT that builds on MSFragger to identify pairs of peptides and their cognate RNA sequences of up to five nucleotides.

## Results

### Non-crosslinked RNA impairs detection of peptide-RNA crosslinks

Current approaches for isolating peptide-RNA crosslinks for LC-MS analysis generally involve in-cell photocrosslinking, protein digestion, RNA enrichment and digestion, and final removal of non-crosslinked RNA (Figure 1A). To investigate why HF-based workflows yielded more peptide-RNA crosslink identifications than nuclease-based methods, we compared their published MS datasets. In representative MS runs from each study, nuclease-treated samples contained abundant adenine- and guanine-derived ions in their averaged MS2 spectra, whereas HF-treated samples did not (Figure 1B). Adenine and guanine ions were also prominent in spectra assigned to peptides crosslinked only to uridine (Figure S1A) and even to non-crosslinked peptides (Figure S1B and S1C). At intact mass level (MS1), ions corresponding to masses of non-crosslinked RNA fragments accounted for a lot of the total intensity (Figure S1D). Together, these observations suggested that highly abundant free RNA fragments, which were removed by chemical digestion with HF but not by nuclease digestion, interfered with peptide identification.

**Figure 1:**
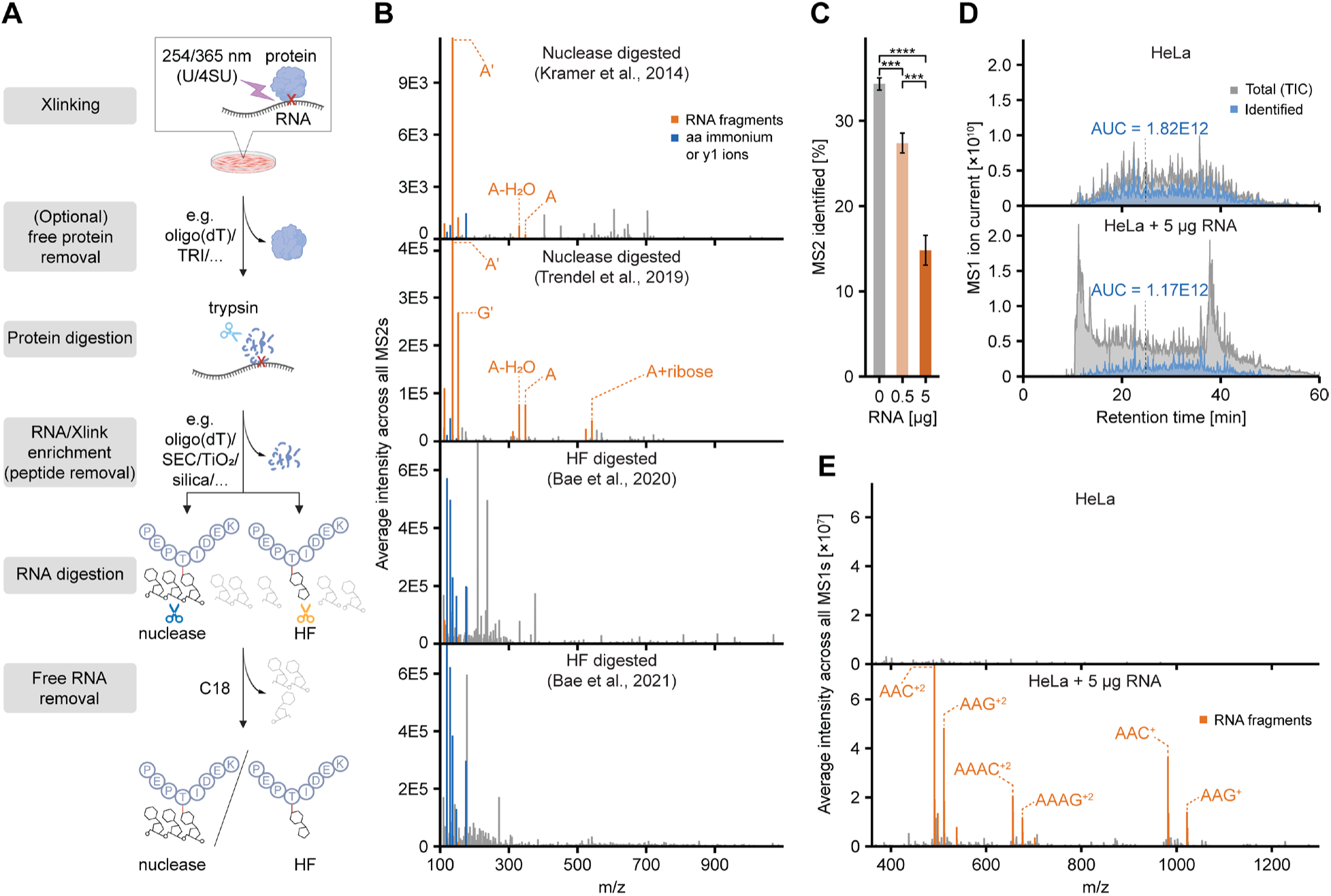
Non-crosslinked RNA contamination in peptide-RNA photocrosslinks impairs MS analysis. **A.** Generalized experimental workflow summarizing published methods for isolating peptide-RNA crosslinks for LC-MS analysis. **B.** Averaged MS2 spectra of four representative MS runs from previous studies. Signals at m/z values corresponding to RNA fragments and amino acid immonium or y1 ions are highlighted. **C.** Bar graph of peptide identification rate upon spike-in of nuclease-digested total RNA into a HeLa tryptic peptide standard. Statistical analysis was performed using a two-tailed unpaired Student’s t-test. ***, p < 0.001 and ****, p < 0.0001. Error bars indicate standard deviation across triplicates. **D.** MS1 chromatogram comparing total ion current and integrated intensity of identified peptides of a HeLa peptide standard with or without RNA spike-in. **E.** Averaged MS1 spectra of a HeLa standard with or without RNA spike-in. Signals of highly abundant RNA fragments are highlighted.

We tested this hypothesis by spiking RNase-digested total cellular RNA into a HeLa peptide standard. Indeed, increasing amounts of RNA digest progressively reduced the peptide identification rate from 34% to 15% (Figure 1C, p = 5.9E-5). Free RNA accounted for a large portion of the total ion current and impaired protein identification throughout the chromatographic gradient (Figure S1E and 1D). At the MS1 level, we detected ions corresponding to RNA oligonucleotides (Figure 1E), similar to those seen in published datasets of nuclease-digested crosslinks (Figure S1D). These species eluted across the gradient and were repeatedly selected for MS2 scans (Figure S1E and S1H), competing with low-abundant peptide-RNA crosslinks for time during data-dependent acquisition (DDA). Unidentified MS2 spectra were strongly dominated by RNA fragment ions (Figure S1F), and even spectra successfully assigned to HeLa peptides contained these ions, although at lower intensity (Figure S1G). The 136 Th adenine ion was by far the most frequently observed and most intense peak of all MS2 spectra in the spike-in samples (Figure S1I). Direct comparison of pure peptide, pure RNA, and mixed scans confirmed that co-isolated RNA suppressed peptide fragment signals (Figure S1J). Such co-isolation resulted from m/z overlap between tryptic peptides and the highly diverse RNA digestion products, which varied in length and sequence and occurred in multiple charge states (Figure 1E).

### Removal of non-crosslinked RNA enhances detection of peptide-RNA crosslinks

We redesigned our nuclease-based workflow^10^ for the isolation of peptide-RNA crosslinks to remove free RNA more efficiently (Figure 2A, S2A). Specifically, we incorporated strong cation-exchange (SCX) solid phase extraction after nuclease digestion to retain peptide-RNA crosslinks while removing free RNA fragments that persist after conventional C18 cleanup^7,8,10,11^. We also optimized the preceding RNA enrichment step by using multiple rounds of isopropanol precipitation, which more effectively removed non-crosslinked peptides. UV spectroscopy revealed that injection-ready samples contained less than 0.1 μg RNA after SCX cleanup, compared with approximately 6 μg after conventional C18 cleanup (Figure 2B). We conducted LC-MS/MS analysis on samples prepared with and without removal of RNA fragments starting from an equal amount of 4SU-labelled, UV-irradiated MCF7 cells (Figure 2C-E). The optimized workflow effectively removed smearing signals derived from non-crosslinked RNA in the MS1 chromatogram, revealing sharp underlying peaks, many of which could be identified as peptide-RNA crosslinks (Figure 2C). The summed intensity of representative RNA fragment ions fell by 54% (Figure 2D). The number of identified peptide-RNA crosslinks (defined as unique modified peptide sequence linked to a unique RNA mass offset, see Methods) increased from approximately 1,700 to more than 4,600 when free RNA was removed (Figure 2E, Table S1). To improve crosslinking efficiency, we integrated our recently reported UV irradiation device for enhanced photoactivation (UVEN^18^) which dramatically reduced the irradiation time from minutes to seconds and enabled irradiation in the original cell culture medium. Strikingly, one second of UVEN irradiation achieved crosslinking comparable to 100 seconds with conventional UV bulbs (Figure S2B). Applying rapid photoactivation combined with improved peptide and RNA removal, we identified 8,725 RNA crosslinks at 2,533 crosslinking sites in 250 proteins in a single MS run, exceeding the yield of the leading HF-based method in all metrics (Figure 2F-2H, Figure S2C, Table S2). This comparison is not strictly matched, as Bae *et al.* analysed mRNA-binding proteins whereas we profiled the total RNA-binding proteome, though comparable crosslinking site numbers were reported for both preparations^16^. Unlike HF digestion, the workflow preserved RNA moieties of one to three nucleotides (Figure 2I). We named this optimized method ‘peptide-RNA crosslink isolation for sequencing by MS’ or pepR-MS.

**Figure 2:**
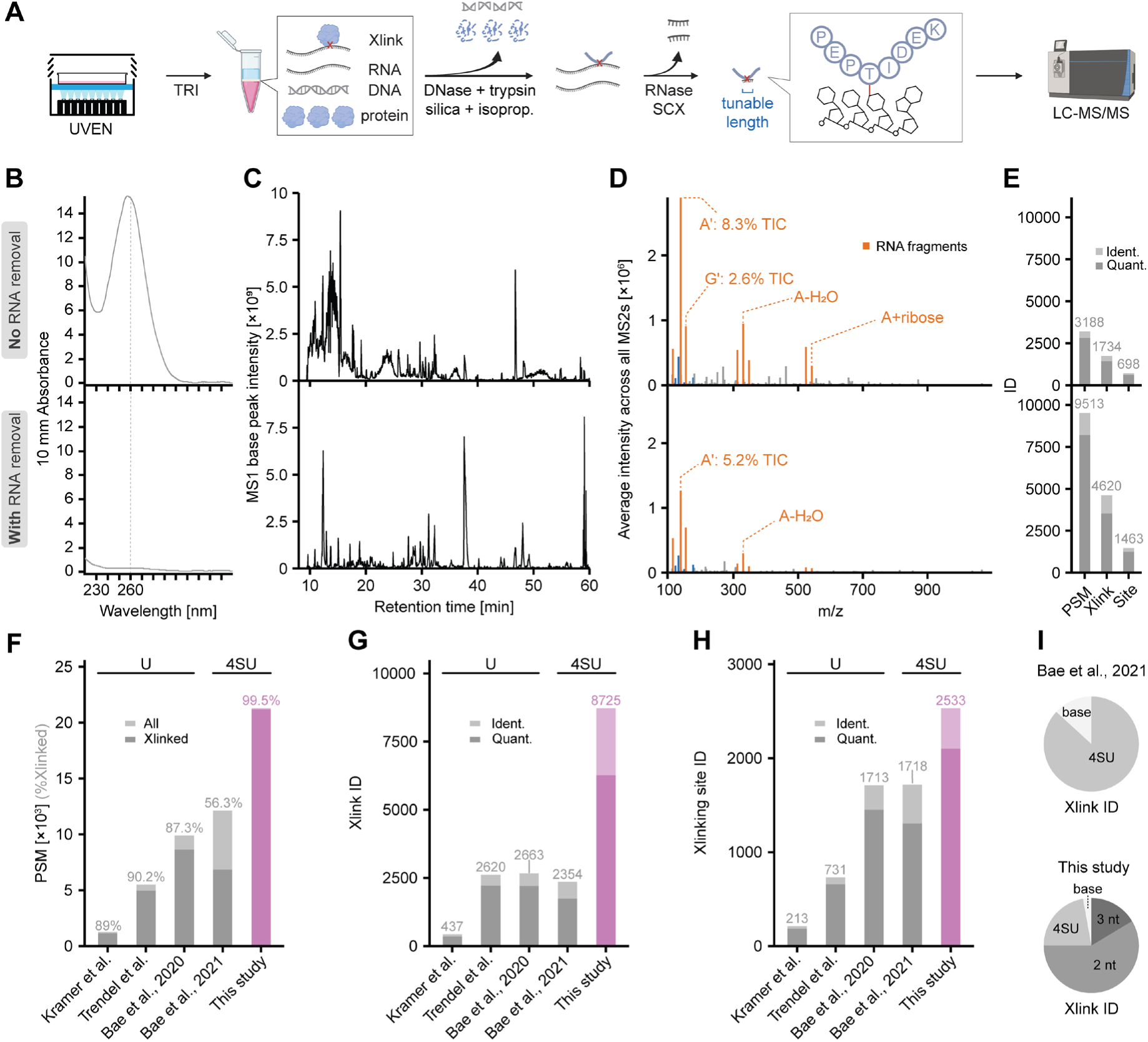
PepR-MS identifies peptide-RNA crosslinks with enhanced robustness and sensitivity. **A.** Experimental scheme of the pepR-MS workflow. **B-E.** Comparison of LC-MS/MS results with and without removal of free RNA. **B.** Line graphs comparing RNA content of the injection-ready MS sample measured by UV spectroscopy. **C.** MS1 base peak chromatograms. **D.** Averaged MS2 spectra comparing fragment ion intensities across all MS2s. Signals of known RNA fragments are highlighted. **E.** Bar graphs comparing identifications by MSFragger using RNA mass offsets up to three nucleotides. **F-H.** Bar graphs comparing published studies with this study. Note that each study either used 254 nm (main crosslinking base U) or 365 nm crosslinking (main crosslinking base 4SU). All data were re-searched with MSFragger (see Methods). **F.** Bar graph showing all PSMs (crosslinked and non-crosslinked). **G.** Bar graph showing identified and quantified crosslinks. **H.** Same as in G for crosslinking sites. **I.** Distribution of RNA length using HF digestion^17^ compared with pepR-MS using nuclease digestion.

### RNA crosslinking sites reveal residue and domain level preferences

To maximize peptide-RNA crosslink coverage in a single cell line, we applied pepR-MS to nuclear and cytosolic fractions of 4SU-labelled MCF7 cells and analysed them separately on an Orbitrap Astral mass spectrometer. Combining pepR-MS with nuclease P1 digestion, we found robust generation of peptide-RNA crosslinks with homogeneous 2 nt RNA chains (91.5% of all crosslinks). Across both fractions, we identified 22,361 unique crosslinks in 468 proteins, providing a rich resource to characterize crosslinking preferences at the protein-RNA interface. To improve crosslink localization at specific sites, we pooled peptide-spectrum matches (PSMs) at the same amino acid position of a protein and defined the best supported site as the focused RNA crosslinking site (hereafter crosslinking site, Figure S3A). Each crosslinking site was supported by a median of seven PSMs, and 41% were covered by more than one peptide sequence. We detected all 20 amino acids at crosslinking sites, with a clear preference for histidine, tyrosine, phenylalanine and glycine compared to the overall amino acid composition of the crosslinked proteins (Figure 3A and 3B, Table S3). This pattern agreed with the preference of 4SU for π-stacking aromatic residues^19^ and was also in line with previous reports^16,17^, except for low crosslinking at cysteine, which will be discussed below. Most crosslinking sites mapped to canonical RNA-binding domains of known RNA-binding proteins, primarily the RNA recognition motif (RRM) and the K homology domain (KH) (Figure 3C and S3B, Table S3). The amino acid frequency at the crosslinking site varied notably between disordered regions and different types of RNA-binding domains (Figure 3D). The RRM was the most frequently crosslinked domain, with 612 unique crosslinking sites mapping to 142 different RRMs from 90 proteins. Multiple sequence alignment of these RRMs revealed 4SU crosslinking hotspots on the central beta sheet formed by β1, β2 and β3 (Figure 3E). To obtain a three-dimensional view, we predicted the structure of a generic RRM based on the consensus sequence of the Pfam profile in complex with the oligonucleotide CAUC using AlphaFold 3^20^. The resulting model showed high prediction confidence (average pLDDT = 91) and resembled experimentally determined RRM-RNA complexes such as the NMR solution structure of the SRSF3 RRM-CAUC^34^ (Figure 3F and S3D). Projecting the crosslinking frequency onto the predicted model indicated strong crosslinking preference around the F/Y residues of the conserved RNP motifs^21^, consistent with experimentally determined RRM-RNA interfaces^22–26^. Apart from canonical RNA-binding domains, more than 400 crosslinking sites localized to IDRs and 45 proteins were crosslinked exclusively within their IDR (Figure S3C). However, crosslinks within IDRs accounted for only 7.4 % of total crosslink intensity and 17 % of PSMs (Figure 3G and S3E). In contrast, the ∼50 % of sites within canonical RNA-binding domains contributed over 75% of total intensity (Figure 3G). This disparity may reflect a longer residence time of RNA contacts within RNA-binding domains compared to more distributed and dynamic interactions within IDRs, which will be further discussed below.

**Figure 3.**
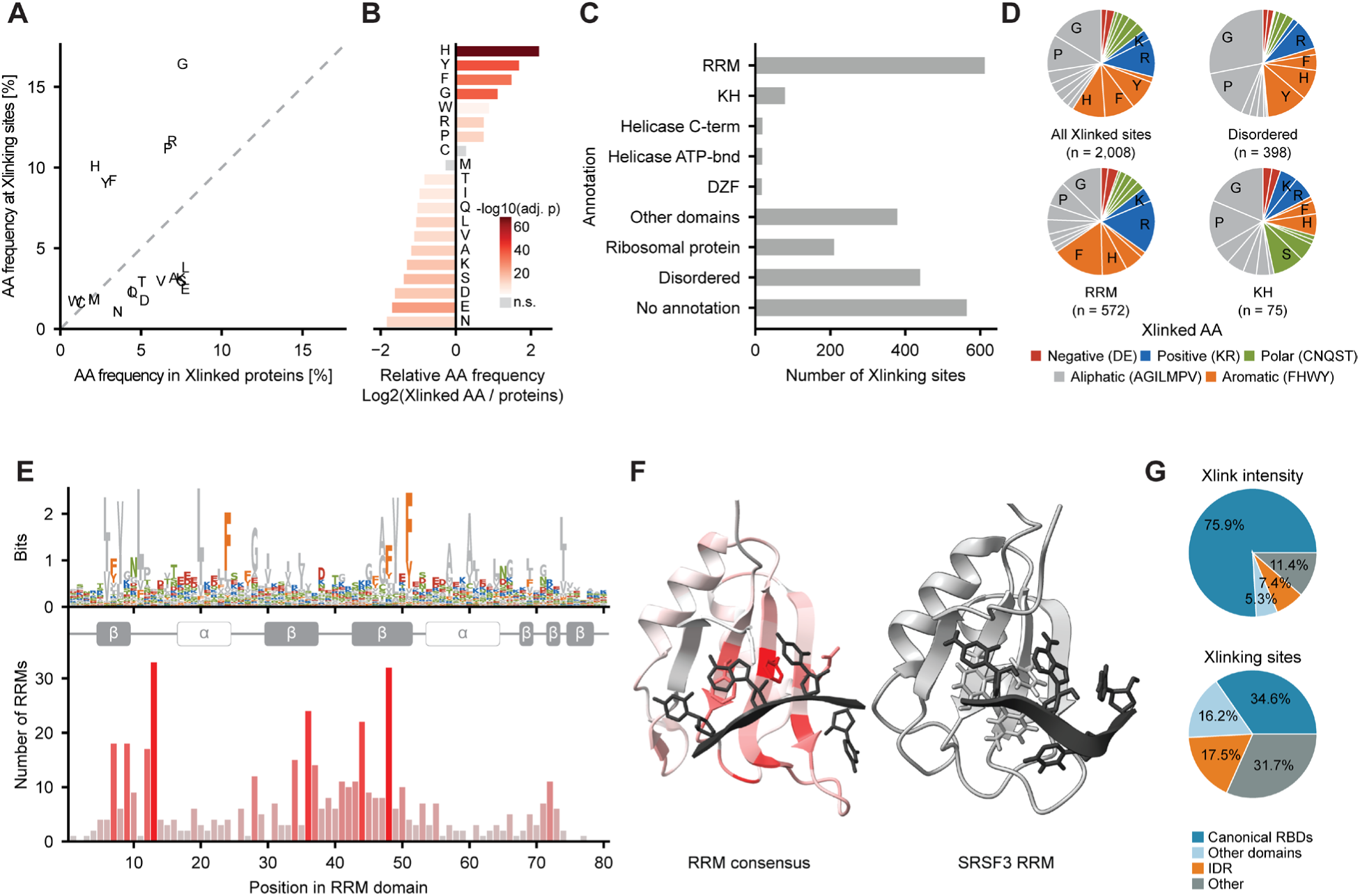
RNA crosslinking sites reveal residue and domain level preferences. **A.** Scatter plot comparing amino acid frequencies in crosslinked proteins with those at crosslinking sites. **B.** Bar plot displaying amino acid frequencies from panel A relative to each other. Colour indicates p-values from Fisher’s exact test after Benjamini-Hochberg correction. **C.** Bar graph showing protein features at crosslinking sites. **D.** Pie diagrams showing amino acid occurrence at crosslinking sites across all proteins, disordered regions, and the two most common RNA-binding domains, sorted by amino acid groups. **E.** Crosslinking sites across all RNA recognition motif (RRM) domains aligned to the RRM Pfam domain definition (top HMM profile). Bars and red colour represent the number of different RRMs found crosslinked at each position. **F.** Left: AlphaFold 3 structure of RRM consensus sequence from Pfam domain definition in complex with CAUC. Red colour indicates the number of RRMs crosslinked shown in panel E. Right: Reference NMR solution structure of the SRSF3 RRM in complex with RNA CAUC (PDB 2I2Y). **G.** Pie charts showing the fraction of crosslink intensity and number of crosslinking sites contributed by crosslinks in canonical RNA-binding domains (RRM, KH, CSD, DZF, Zinc finger, Helicase C-term, DEAD/DEAH box helicase, and dsRBD), other InterPro domains, IDRs and other regions.

### Crosslinks in IDRs show local sequence features absent from structured domains

In molecular condensates, RNA has been shown to promote aggregation, e.g. via scaffolding in paraspeckles^27,28^ or nucleoli^29^, but in other cases to promote condensate dissolution, e.g. in transcriptional bursting^30,31^. Where RNA interactions occur in IDRs and how they contribute to attractive or repulsive behaviour remain incompletely understood. Visually inspecting the positions of over 400 crosslinking sites in IDRs identified by pepR-MS, we noticed that they clustered within restricted loci, with a median nearest-neighbour distance of three amino acids. This spacing was smaller than expected for randomly distributed sites (p < 1E-4, Figure S4A), suggesting that the captured crosslinks reflect defined contact regions rather than arbitrary contacts along the IDR. Moreover, crosslinking sites often overlapped with disordered sequence motifs known to bind nucleic acids. In SFPQ, for example, crosslinks clustered at an RG/RGG box and at aromatic, G-rich stretches (Figure 4A). To better understand the molecular grammar of the RNA-interacting IDRs, we compared their classification by NARDINI+/GIN which clusters human IDRs into common classes based on amino acid compositional and patterning features^32^. Crosslinked IDRs were significantly enriched in blocks of G and polar amino acids, high R fraction, as well as high aromatic fraction, especially Y (Figure 4B and Figure S4B). They were depleted in clusters of negatively charged residues consistent with electrostatic complementarity to the phosphate backbone^33^.

**Figure 4.**
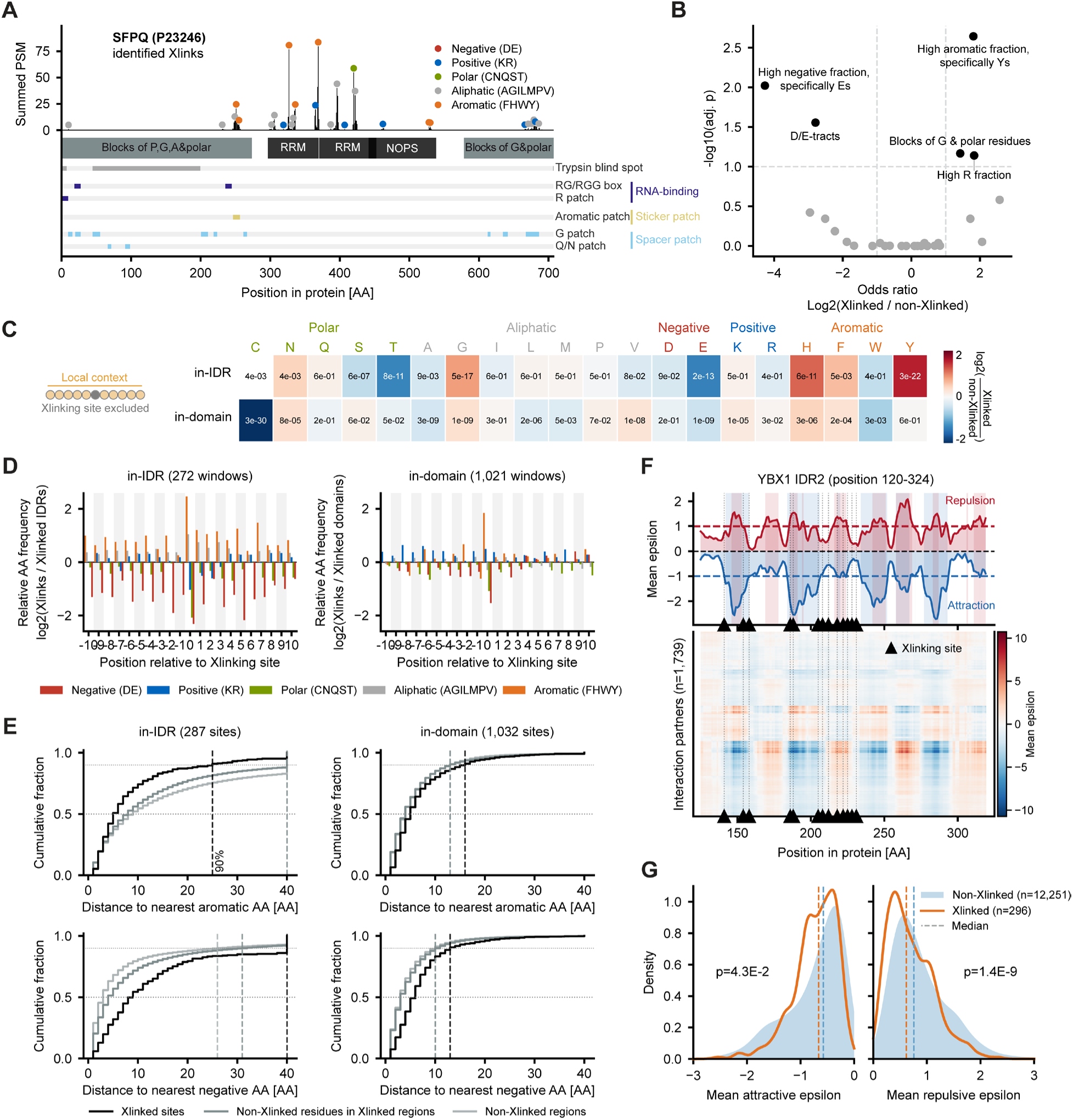
RNA crosslinking sites show IDR-specific local sequence features absent from structured domains. **A.** Summed PSM evidence per residue along SFPQ, with crosslinking sites coloured by residue. Structured domains and GIN-cluster-annotated IDRs, together with RNA-binding motifs, sticker/spacer patches and tryptic blind spots are annotated. **B.** Volcano plot showing GIN-cluster enrichment or depletion in crosslinked vs. non-crosslinked IDRs. P-values were calculated using two-sided Fisher’s exact tests, with the Benjamini-Hochberg correction across all clusters. Significant enrichment and depletion are highlighted. **C.** Heatmap showing amino acid enrichment or depletion within five residues on either side of crosslinking sites, relative to non-crosslinked residues in crosslinked IDRs (top) or domains (bottom). Colours indicate log2 fold-change in AA fraction; cells are annotated with p-values calculated using a two-sided Mann-Whitney U test with Benjamini-Hochberg correction. **D.** Normalized sequence logos showing positional features within ±10 residues of crosslinking sites. Enrichment and depletion are shown as log2 fold change relative to the amino acid composition of IDRs or domains in crosslinked proteins. **E.** Cumulative distribution plots showing distances to the nearest aromatic (top) or negatively charged residue (bottom) in IDRs (left) and domains (right). Crosslinking sites are compared with non-crosslinked residues in crosslinked and non-crosslinked regions. **F.** FINCHES-predicted residue-resolved attractive and repulsive interactions along a YBX1 IDR, averaged across all partners (top) or shown per partner (bottom). Crosslinking sites identified in this IDR are annotated. **G.** Distribution of per-residue mean interaction propensity for non-crosslinked and crosslinked residues in crosslinked IDRs. Negative values indicate attraction; positive values indicate repulsion. See Methods for analysis and statistical procedures.

We next asked whether these sequence features were global properties of the crosslinked IDRs or locally concentrated around RNA crosslinking sites. In IDRs, a window of ten residues around the crosslinking site (but excluding the crosslinked residue itself) showed characteristic amino acid enrichments and depletions that differed from those around sites in structured domains (Figure 4C). Aromatic residues, especially Y, were strongly enriched locally. Histidine was also enriched near crosslinks despite showing no global enrichment in crosslinked IDRs (Figure 4C and S4B). Aromatic residues were otherwise rare in both non-crosslinked IDRs and regions of crosslinked IDRs outside the local contact site (Figure S4C). This pattern is consistent with evidence that F and Y can be essential for IDR-RNA binding, as in the F/YGG motif of hnRNPA2^34^. G was also enriched near crosslinks and may provide the flexibility required to position aromatic or R side chains for RNA contact^35,36^. While globally enriched in crosslinked IDRs, R was not enriched in the direct neighbourhood of crosslinking sites, potentially because trypsin produced short peptides from K/R-rich sequences that escaped our LC-MS detection. Acidic residues, especially E, were more strongly depleted around crosslinking sites than their global depletion across the entire crosslinked IDRs (Figure S4C). In contrast to other studies^16,17^, C showed no enrichment at crosslinking sites (Figure 3B) and was strongly depleted around crosslinking sites in structured domains (Figure 4C). Structural analysis has shown that the previously reported RNA-C crosslinks often lie outside structurally resolved protein-RNA interfaces^37^. However, why they were detected so abundantly in previous studies and were absent from our data remains unclear to us.

As expected, sequence logos around IDR crosslinks showed little positional conservation (Figure S4D). Aromatic residues were enriched and acidic residues depleted uniformly throughout five amino acids on each side of the crosslinking sites, without a clear positional motif (Figure 4D). This local pattern was specific to crosslinked IDRs, where 90% of crosslinked residues had an aromatic residue within 25 amino acids, whereas the corresponding distance was larger 40 amino acids for non-crosslinked residues both in the same IDRs and in IDRs without crosslinks (Figure 4E). Thus, unlike the residue-focused contacts in structured domains, RNA contacts in IDRs extend across short aromatic and G-rich segments depleted of D and E, and enriched in Y, H and F. Aromatic residues can act as stickers mediating IDR-IDR interactions in the stickers-and-spacers model of condensate assembly^38^. Indeed, crosslinks were significantly closer to aromatic and G-rich patches, consistent with the local sequence analysis (Figure S4E). Because RNA-binding proteins frequently interact with one another^39^, we used FINCHES^40^ to predict interactions between each crosslinked IDR and 1,739 IDRs from 791 RNA-binding proteins expressed in MCF7 cells. Figure 4F shows on the example of YBX1 that even within the same IDR, RNA interaction occurred in stickers as well as spacers. Across all crosslinked IDRs, however, crosslinking sites tended to occur in regions predicted to mediate attractive rather than repulsive IDR interactions (Figure 4G).

### Sequential sequencing of peptide and RNA in crosslinks

Peptide-RNA crosslinks with sufficient amino acid and nucleotide chain length could locate a protein-RNA interface on both the protein and RNA side. While unambiguous mapping to the transcriptome would require a median RNA length of 14 nucleotides^41^, shorter sequences could already provide insight into the binding preferences of individual RNA-binding domains. These typically interact with degenerate motifs of 3-8 nucleotides^42,43^. We therefore tested whether nuclease choice could tune the retained RNA length. Nuclease P1, RNase A, and a nuclease mixture produced mainly two-nucleotide adducts; RNase T1 produced mainly two- to three-nucleotide adducts; and benzonase shifted the fraction towards three- to five-nucleotide adducts (Figure 5A and Table S4). The mass offsets detected by MSFragger aligned with the known cleavage specificity of each nuclease (Figure S5A).

**Figure 5:**
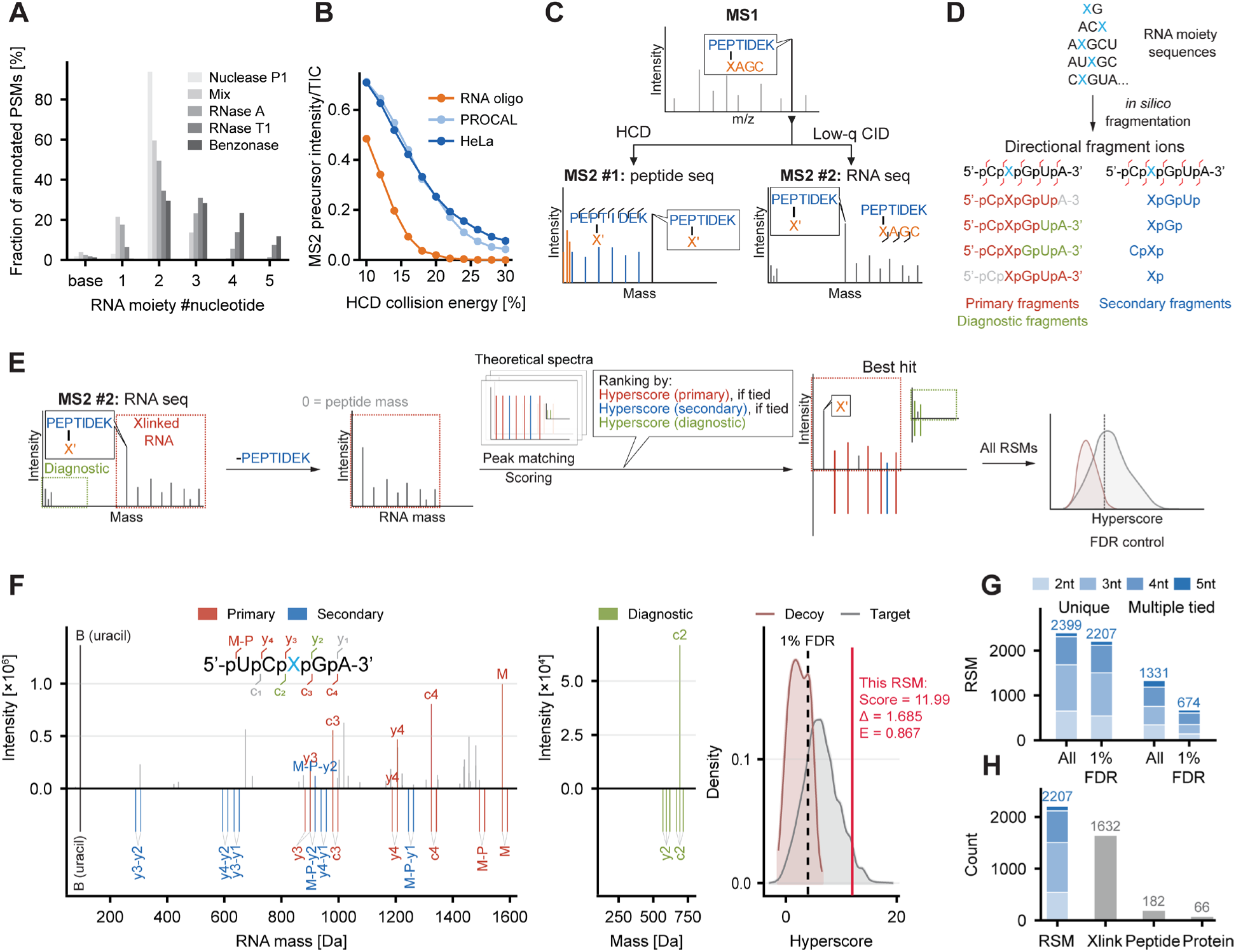
Sequencing peptide and RNA moieties of peptide-RNA crosslinks in the same MS run. **A.** Histogram of PSMs with RNA moieties of different lengths identified by MSFragger offset search on samples using various nuclease combinations. **B.** Scatter plot illustrating the fragmentation behaviour of the synthetic RNA oligo UAGUAG, a HeLa peptide standard, and the synthetic peptide standard PROCAL across increasing HCD collision energies. The mean ratio of precursor intensity to total MS2 ion current is shown. **C.** Schematic of the MS method for sequential sequencing of peptide-RNA crosslinks. **D.** Schematic of theoretical spectrum database generation. **E.** Schematic of the SALT pipeline for sequencing RNA moieties of peptide-RNA crosslinks. **F.** Example RSM identifying the RNA moiety 5’-UCXGA-3’ (X, crosslinked 4SU) crosslinked to peptide ITLSKhQNVQLPR from PTBP1. Left, charge deconvoluted, peptide-mass-subtracted experimental RNA spectrum (top) matched to theoretical primary and secondary fragments (bottom). Middle, match of diagnostic ions for the same RSM using the charge-deconvoluted but non-peptide-mass-subtracted experimental spectrum. Right, Hyperscore distribution of target and decoy matches from the same MS run. The Hyperscore threshold at 1% FDR is indicated by a dashed line and the Hyperscore of this RSM by a red line. Δ, difference in Hyperscore between the highest and the second highest ranked candidate sequences. E, Expectation-value of the best match. **G.** Bar plot showing the number of RSMs assigned to a unique ‘winner’ sequence or multiple ‘tied’ sequences for RNA lengths of 2-5 nt, before and after filtering at 1% FDR. **H.** Bar plot showing unique RSM, crosslink, peptide and protein IDs of one MS run.

Mass offsets only reveal the composition of the crosslinked RNA but not its sequence. To determine both RNA and peptide sequences, we first compared their fragmentation under the LC-MS conditions used for crosslink analysis. The synthetic RNA UAGUAG fragmented at substantially lower collision energies than either a HeLa peptide standard or synthetic PROCAL peptides^44^ (Figure 5B), consistent with previous analyses of peptide-oligonucleotide conjugates under positive ESI-MS conditions^45^. This offered a way to decouple sequencing of the crosslinked RNA and peptide moieties. We implemented an acquisition method that recorded two consecutive MS2 scans from the same precursor, using higher-energy collisional dissociation (HCD) to fragment the peptide and low-q ion-trap collision-induced dissociation (CID) to fragment the RNA (Figure 5C). Ion-trap CID is widely used for tandem MS of oligonucleotides^46,47^, and recent work has shown that reducing the Mathieu equation q value during activation limits secondary fragmentation^48–50^. We applied this strategy to benzonase-digested crosslinks containing RNA adducts of up to five nucleotides (Figure 5A). Following acquisition, MSFragger was then used to identify MS2 spectra of peptides with mass offsets corresponding to RNA chains of up to five nucleotides (Figure S5B). This identified 4,509 MS2 spectra for RNA-crosslinked peptides from the HCD scans and 1,326 MS2 spectra from the low-q CID scans, most of which overlapped with HCD identifications (Figure S5C).

Sequencing the RNA moiety from the low-q CID spectra required the development of a dedicated computational pipeline, as no current software supports sequencing of the RNA moiety in peptide-RNA crosslinks in positive-ion mode^51–53^. We therefore developed the analysis extension SALT (Sequential Analysis of peptide-xLinked nucleoTides) to identify RNA sequences from our dual-MS2 data after MSFragger peptide assignment. For RNA chains generated by nonspecific nuclease digestion (e.g., benzonase), any nucleotide sequence carrying a crosslinking U at any position can occur in principle. We therefore generated a sequence database of all 3,710 possible RNA sequences up to five nucleotides and computed theoretical spectra based on expected CID fragment ions. This included intact precursors, peptide-crosslinked primary and secondary fragments, and peptide-free terminal diagnostic fragments of at least two nucleotides (Figure 5D; see Methods). For each low-q CID scan, SALT subtracts the unmodified peptide mass assigned by MSFragger in the HCD spectrum and matches the remaining RNA peaks to the theoretical RNA library. A decoy library of chemically modified RNA species provides false-discovery-rate (FDR) control (Figure 5E; see Methods). Figure 5F shows an example RNA-spectrum-match (RSM) identifying the 5-nt RNA 5’-UCXGA-3’ crosslinked to a PTBP1 peptide. This RNA sequence showed the highest Hyperscore among 1,398 candidate RNA sequences.

Of 4,509 low-q CID scans with paired peptide identifications, 3,730 matched at least one theoretical RNA spectrum based on one or more matched fragment ions. Approximately one-third of identifications initially produced tied scores because incomplete fragmentation ladders could support several sequences, despite consideration of secondary and diagnostic peak matches. However, most of the ambiguous matches were removed at a 1% FDR threshold, confirming that FDR thresholding did remove low confidence identifications (Figure 5G). RSMs with more than two nucleotides were more likely to pass the FDR filter because their spectra provided more informative fragment ions. Consequently, the longest RNA chains were identified with the greatest confidence. In one LC-MS run, SALT identified more than 1,600 unique crosslinks from 66 proteins with both peptide and RNA sequences (Figure 5H, Table S5).

### Peptide-RNA crosslinks capture diverse interfaces in ribonucleoprotein complexes

We validated peptide-RNA crosslinks obtained by our approach using the crystal and cryo-EM structures of the human spliceosome (Figure 6A) and the ribosome (Figure 6B-E and Figure S6). Twelve unique ribosomal crosslinks were identified with RNA moieties longer than two nucleotides represented by 33 RSMs, of which 29 RSMs had a distance smaller than 20 Å between the amino acid Cα and uridine N1 at their corresponding positions in the ribosome structure (Figure S6A). Nine out of twelve crosslinks had a Cα-N1 distance below 10 Å, similar to the distance range observed in recent UV-crosslinking studies^54,55^.

**Figure 6.**
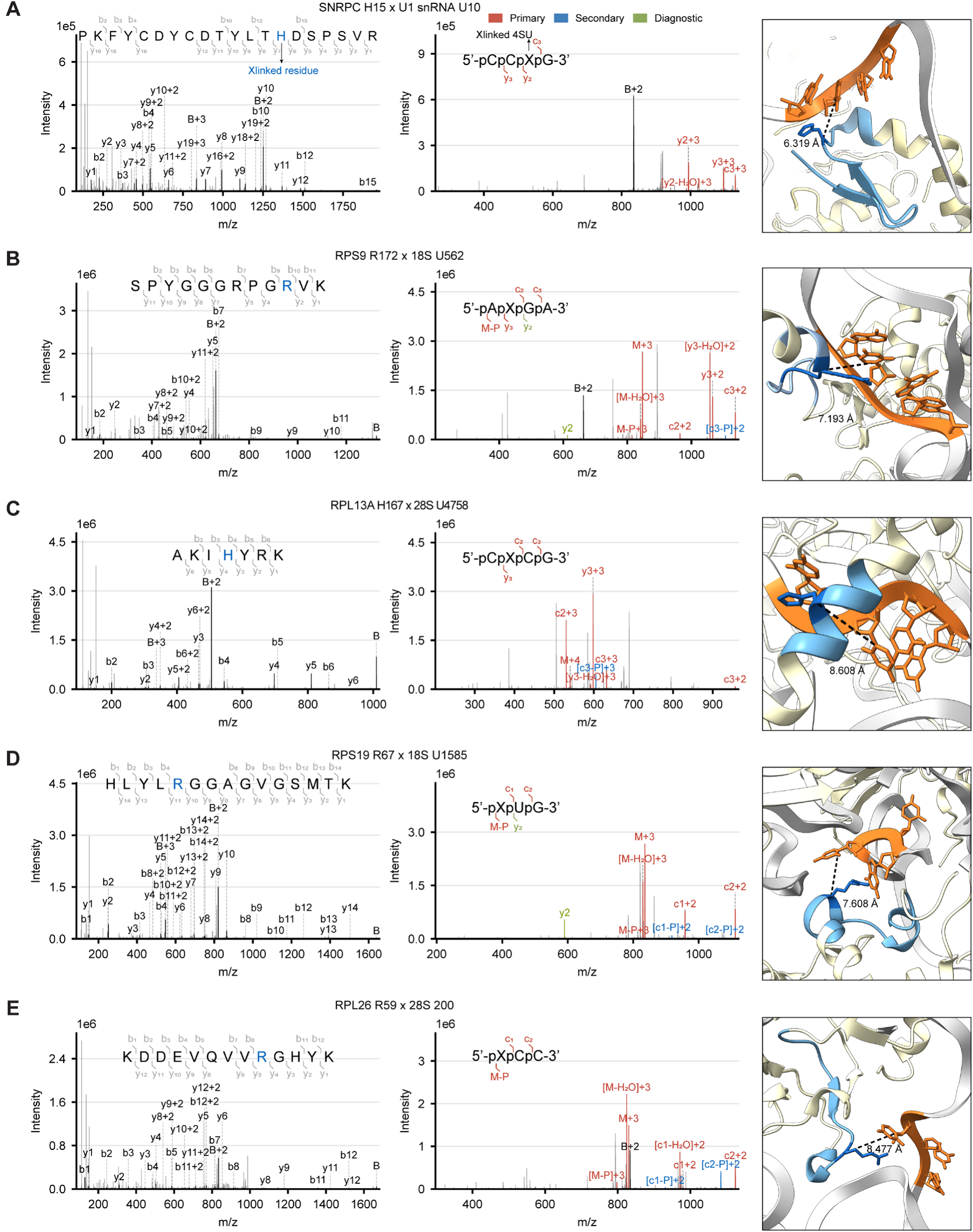
Protein-RNA crosslinks capture diverse interfaces in ribonucleoprotein complexes. **A.-E.** Left and middle panels: pairs of MS2 spectra (left: HCD, middle: low-q CID) for sequential sequencing of ribosomal and spliceosomal crosslinks. Crosslinked residues are highlighted in blue. If more than one residue is equally likely to be crosslinked, the closest residue to the identified 4SU in the structure is highlighted. X, crosslinked 4SU; B, uracil (base)-crosslinked peptide; M, precursor with intact RNA chain. Right panel: mapping of sequenced crosslinks to the crystal structure of human U1 hnRNP (panel A, PDB 4PJO) and the cryo-EM structure of human 80S ribosome (panels B-E, PDB 4UG0). Light blue: identified peptide; dark blue: crosslinking site(s); orange: crosslinked RNA segment. The distance between the Cα of the crosslinked amino acid residue and the N1 of crosslinked 4SU is annotated.

Crosslinks were identified from diverse structural contexts. For example, R17 and R58 of RPS9 crosslinked to the first three nucleotides at the 5’ end of 18S rRNA, positioned on opposite sides of the RNA in the cryo-EM structure (Figure S6B and S6D). R17 lies in an unstructured N-terminal coil, whereas R58 lies in the folded core (Figure S6B). R172, crosslinked to positions 561-564 of 18S rRNA, is located in the low-complexity C-terminal region of the protein (Figure 6B and S6B). We also found RPL13A crosslinked to the 28S rRNA via its C-terminal sequence (Figure 6C). RPL13A has been shown to be phosphorylated and released from the ribosome upon prolonged interferon-γ treatment and participates in GAIT-mediated translational silencing by binding GAIT elements in the 3′ UTRs of specific mRNAs^56^. Crosslinked H167 is located immediately adjacent to residues R169-K171, which are dispensable for ribosome incorporation but required for GAIT-mediated translational silencing^57^. In RPS19, crosslinks at Y48 and R67 flank the central α-helix spanning residues 52-67 (Figure 6D and S6E), a hotspot for missense mutations in Diamond-Blackfan anaemia (DBA)^58^ (Figure S6C). Deletion of RPS19 residue 67 has recently been used to establish an experimental DBA model^59^, suggesting the potential of pepR-MS to investigate how disease-associated mutations alter protein-RNA interfaces in the ribosome. Indeed, we detected another crosslink at R59 of RPL26 within a segment lost in a DBA-associated frameshift mutation, which causes defective pre-rRNA processing in patient cells^60^ (Figure 6E). These examples show that peptide-RNA crosslinks with longer RNA chains identified by pepR-SALT could be validated against published structures of protein-RNA complexes and mapped several functionally relevant regions associated with congenital ribosomopathies.

## Discussion

MS analysis of peptide-RNA crosslinks can resolve protein-RNA interfaces to single amino acids in living cells. However, its broader application has been limited by several technical challenges. First, the low abundance of peptide-RNA crosslinks due to the low efficiency of UV crosslinking and the large excess of non-crosslinked peptides requires highly efficient enrichment methods to recover pure crosslinked species^7,54,61^. Second, RNA-chain complexity has been assumed to complicate MS/MS spectra and expand the MS search space, impairing interpretation of LC-MS/MS data^12–14,16,17^.

We identify free RNA contamination, rather than the size of the MS data search space, as the main obstacle to detecting peptide-RNA conjugates by MS. By removing non-crosslinked RNA fragments while retaining peptide-RNA crosslinks, even with longer RNA chains, pepR-MS led to four distinct improvements. First, the much lower free RNA content of the injection-ready MS sample led to much larger amounts of peptide-RNA crosslinks that could be loaded onto the LC system without saturating it. Second, sample purity strongly decreased carry-over between LC-MS runs allowing for many consecutive injections without compromising LC performance. Third, pure preparations allowed for a wider dynamic range of crosslink detection, improving sensitivity for low-abundant species, which were previously masked by RNA fragments. Fourth, pervasive MS2 co-isolation of free RNA fragments was effectively reduced, allowing peptide fragment signals to be detected at higher intensities and improving MS2 spectral quality.

A limitation of our and previous approaches, particularly in the analysis of RNA-IDR contacts, is that K/R-rich sequences are poorly represented when using trypsin digestion. Proteases such as Glu-C may therefore reveal contacts underrepresented in the current dataset. Conversely, we find aromatic IDR-RNA interactions very well represented in our data. As aromatic amino acids have the greatest photochemical reactivity towards nucleic acids^19,37^, it remains unclear to what extent their frequent detection reflects genuine IDR-RNA contacts in cells rather than an intrinsic crosslinking bias. Because most crosslinks actually occur with chemically rather inert glycine, and because aromatic amino acids are essential components of known disordered RNA-binding motifs (e.g. F/YGG^34^), we think that their enrichment at crosslinking sites reflects increased RNA residence time at this part of the IDR. In how far this implies RNA binding, increased interactions within molecular condensates, or bystander crosslinking, requires validation on a case-by-case basis.

PepR-MS also allows the length of the retained RNA to be tuned through nuclease choice, which enabled us to sequence both components of peptide-RNA crosslinks bearing RNA chains of up to 5 nt in length. While this suggests that proteome-transcriptome-wide mapping of protein-RNA interactions might in principle be possible, it also reveals the challenges yet to be overcome. Unambiguous assignment of RNA moieties to their transcripts of origin requires crosslinks with longer RNA sequences. Shorter nuclease digestion times or physical shearing methods such as sonication may preserve longer RNA. The acquisition capacity of current MS instrumentation will remain a constraint because of the enormous number of possible peptide-RNA sequence combinations. Many biological questions, however, may be addressed without exhaustive coverage of all-against-all but only identifying the predominant RNA partners of several hundred proteins. The complexity of the interaction space can also be reduced experimentally by enriching specific cellular fractions, ribonucleoprotein complexes, or individual proteins before MS analysis, as in CLIP-seq workflows^4,62^. While measuring longer RNA chains becomes more challenging in terms of both sample complexity and MS methodology, we expect database searching to become more straightforward. Longer RNA sequences carry more information and can therefore be identified with greater confidence. Established software solutions for RNA sequencing by MS could be repurposed for this application^51–53^. PepR-SALT provides a complete workflow for site-specific mapping of protein-RNA interactions within a single LC-MS experiment and offers a route forward to proteome- and transcriptome-wide characterization of protein-RNA interactions by LC-MS at single-amino acid and nucleotide resolution.

## Methods

### Cell culture

MCF7 cells (human, female breast adenocarcinoma) were obtained from ATCC. Cells were cultured in Dulbecco’s Modified Eagle’s Medium (DMEM, PAN-Biotech GmbH) supplemented with 10% fetal bovine serum (FBS, PAN-Biotech GmbH), in a 37°C incubator with a CO_2_ concentration of 5%. The cells were seeded in 150-mm tissue culture dishes (Sarstedt) with 20 mL culture medium at 2 million cells per dish. After three days of culture, cells were labelled for 18 h under original cell culture conditions with 100 µM 4SU (Biosynth) by adding from a 100 mM stock in DMSO directly to the culture medium.

### RNA digest spike-in experiments

RNA was extracted from 4SU-labelled MCF7 cells with TRI reagent (or TRIzol, a mixture of guanidine thiocyanate and phenol) (Sigma Aldrich) according to the manufacturer’s instructions. After isopropanol precipitation, RNA pellets were dissolved in 50 mM Tris-HCl (pH 7.5) and adjusted to a concentration of 1 µg/µL, as determined by NanoDrop UV spectroscopy (Thermo Fisher Scientific). For 100 µg RNA (in 100 µL), residual proteins were digested by adding 20 µL proteinase K (NEB) at 20 µg/µL and incubating at 55°C for 15 min. The sample was cooled to room temperature, then 400 µL RNA Lysis Buffer (Zymo research, Quick-RNA Miniprep Kit) and 500 µL ethanol were added, mixed by inversion, purified over spin columns according to the manufacturer’s instructions and eluted with three times 100 µL ultrapure water. To 250 µL eluate, 5 µL 500 mM ammonium bicarbonate (final 10 mM), 2 µL RNase A (0.5 µg/µL, Thermo Fisher Scientific) and 2 µL RNase T1 (0.5 µg/µL, Sigma-Aldrich) were added and incubated at 37°C overnight. After RNA digestion, 0 (vehicle control), 0.5 or 5 µg RNA fragments as determined by NanoDrop were added to 150 ng HeLa peptide standard (in-house) in triplicate and the samples were lyophilized in a SpeedVac (Thermo Fisher Scientific) for LC-MS/MS analysis. The 150 ng HeLa peptide input corresponds to the routine QC standard for the nanoflow LC-MS setup used in our laboratory, chosen to match column capacity and MS sensitivity. The amount of spiked in RNA aligns the approximately 6 µg of RNA in C18-desalted crosslink samples (Figure 2B), so that the 5 µg spike-in reproduces the free RNA contamination in practice, with 0.5 µg representing a ten-fold lower load. Wash blanks after both 0.5 µg and 5 µg samples showed intense RNA-derived signal and the trap column had to be replaced after three replicates (raw files were deposited to MassIVE).

### UV crosslinking

For bulb-based UV crosslinking, culture medium was removed from 4SU-labelled MCF7 cells, which were washed with 10 mL ice-cold Dulbecco’s phosphate buffered saline (PBS, with calcium chloride and magnesium chloride, Sigma-Aldrich), covered with 20 mL ice-cold PBS and kept on ice during irradiation. Photocrosslinking was performed in a conventional UV crosslinker (Biolink, Vilber) with 365 nm bulbs for 5, 10, 100 or 500 s (for UV irradiation timeline samples, Figure S2B) or to a total irradiation energy of 200 mJ/cm^10^ (∼50 s, for workflow comparison samples, Figures 2B-2E). Cells were scraped into the ice-cold PBS they were irradiated in and pelleted in 3-dish batches by centrifugation at 500 g for 5 min for further processing.

For LED-based UV crosslinking, cells were transferred directly from the incubator to the high-intensity irradiation device UVEN^18^ and irradiated in their original culture medium with 365 nm LEDs for 1, 3, 5 or 10 s (for UV irradiation timeline samples, Figure S2B) or 5 s (all other peptide-RNA crosslink samples if not specified above). Cells were washed twice with 5 mL ice-cold PBS, scraped into 15 mL ice-cold PBS and pelleted in 3-dish batches by centrifugation at 500 g for 5 min for further processing.

### PepR-MS workflow for peptide-RNA crosslink isolation

Cell pellets from three 150 mm dishes were lysed in 3 mL TRI reagent by pipetting until completely homogeneous. Lysates were combined with 600 µL chloroform, mixed by inversion and centrifuged at 3000 g for 20 min at 4°C. The interphase was transferred to a fresh tube, dissolved in 1 mL TRI reagent by pipetting until completely homogeneous, combined with 200 µL chloroform, mixed by inversion and centrifuged at 12000 g for 10 min at 4°C. The aqueous and organic phases were removed, and the interphase was disintegrated in recovery buffer (10 mM Tris-HCl, 1 mM EDTA and 1% SDS, pH=7.4) until completely dissolved. For every 1 mL of dissolved sample, 100 µL 5 M NaCl and 1 mL isopropanol were added, mixed by inversion and incubated at -20°C overnight to precipitate RNA. Samples were centrifuged at 17000 g for 30 min at -11°C. The supernatant was discarded, and the pellets were washed with 1 mL 70% ethanol and combined into one tube for a 3-dish sample. Residual ethanol was removed to completion. Combined 3-dish pellets were hydrated with 760 µL ultrapure water, detached from the tube wall by gentle pipetting and incubated for 15 min on ice. DNA was digested by adding 90 µL 10× DNase I buffer (NEB), 2 µL RNasin ribonuclease inhibitor (Promega) and 50 µL 2000 units/mL DNase I (NEB), followed by incubation for 1 h at 37°C with shaking at 500 rpm. Protein was digested by adding 5 µL 20% SDS, 5 µL 1 M DTT and 60 µL 1 µg/µL trypsin (Roche) and incubating for 3 h at 37°C with shaking at 500 rpm. For the workflow comparison experiment (Figure 2B-2E), two 3-dish samples were pooled and split equally to minimize technical variation up to this step.

For crosslinking site localization samples (Figure 3 and 4), cell pellets from six 150 mm dishes were fractionated prior to processing. Pellets were lysed in 6 mL nuclei extraction buffer (10 mM Tris-HCl pH 7.5, 60 mM KCl, 1.5 mM MgCl_2_ and 0.1% NP-40) by pipetting on ice until completely homogeneous and centrifuged at 2000 g for 5 min at 4°C. The supernatant (cytosolic fraction) was separated from the pellet (nuclear fraction) and each fraction was processed as a 3-dish equivalent. The nuclear fraction was resuspended in 2 mL TRI reagent and processed as described above. The cytosolic fraction was precipitated directly with isopropanol and processed as described above from the precipitation step onward, except that digestion was performed with 100 µL trypsin (1 µg/µL) rather than 60 µL.

After trypsin digestion, the ‘no RNA removal’ sample in the workflow comparison experiment (upper panel of Figure 2B-2E) was adjusted to a final volume of 1 mL with ultrapure water and directly processed by silica column purification as described below. For all other peptide-RNA crosslink isolations, 100 µL 20% SDS was added to each trypsin-digested 3-dish sample, which was incubated for 15 min at 60°C with shaking at 500 rpm. RNA was precipitated by adding 100 µL 5 M NaCl and 1 mL isopropanol, mixing by inversion and incubating at -20°C for at least 30 min. Precipitated samples were centrifuged at 17000 g for 30 min at -11°C. The supernatant was discarded, and pellets were washed with 1 mL 70% ethanol. For all 3-dish samples (all samples unless specified as 12-dish below), pellets were resuspended in 1 mL ultrapure water. For all 12-dish samples (peptide-RNA crosslink sample for ID comparison in Figure 2F-2I and Figure S2C and tunable-length samples in Figure 5), four 3-dish pellets were combined and resuspended in 2 mL ultrapure water.

To each mL of resuspended sample, 3.5 mL RLT buffer (Qiagen, RNeasy Midi Kit) was added and mixed by inversion. The mixture was incubated at 85°C for 15 min with shaking at 500 rpm and cooled to room temperature before adding 2.5 mL ethanol. Each 3-dish (total volume 7 mL) or 12-dish sample (total volume 14 mL) was loaded onto one RNeasy Midi column in 3.5 mL increments, washed with 8 mL (3-dish samples) or 12 mL (12-dish samples) RW1 buffer and 9 mL RPE buffer, and eluted with 1 mL (3-dish samples) or 6 mL (12-dish samples) ultrapure water. Eluates were heated to 85°C for 1 min with shaking at 500 rpm. RNA was precipitated by adding 100 µL 5 M NaCl and 1 mL isopropanol to each 1 mL eluate, mixing by inversion and incubating at -20°C overnight.

Precipitated samples were centrifuged at 17000 g for 30 min at -11°C. The supernatant was discarded, and the pellets were washed with 1 mL 70% ethanol, then resuspended with 100 µL (3-dish) or 400 µL (12-dish) 50 mM Tris-HCl (pH 7.5) and heated to 85°C for 1 min with shaking at 500 rpm. RNA amounts were determined by NanoDrop. For each 100 µg RNA, 2 µL RNase A (0.5 µg/µL) and 2 µL RNase T1 (0.5 µg/µL) were added and incubated for 16 h at 37°C with shaking at 500 rpm, unless specified otherwise. For crosslinking site localization samples (Figure 3 and 4), each 100-µg RNA was incubated with 1 µL nuclease P1 (100,000 units/mL, New England Biolabs) and 2 µL benzonase (0.1 µg/µL). For tuning the RNA length with different nucleases (Figure 5), all samples were supplemented with 20 µL 100 mM MgCl_2_ (final 5 mM). For these samples, each 100 µg RNA was incubated with a) 1 µL nuclease P1 (100,000 units/mL) and 2 µL benzonase (0.1 µg/µL); b) 2 µL RNase A (0.5 µg/µL), 2 µL RNase T1 (0.5 µg/µL) and 2 µL benzonase (0.1 µg/µL; GENIUS nuclease, Santa Cruz Biotechnology); c) 2 µL RNase A (0.5 µg/µL); d) 2 µL RNase T1 (0.5 µg/µL); and e) 2 µL benzonase (0.1 µg/µL). For dual sequencing samples (Figure 5 and 6), benzonase (same as group e above) was used.

The ‘no RNA removal’ sample in the workflow comparison experiment (upper panel of Figure 2B-2E) was adjusted to 150 µL with water and formic acid to a final concentration of 0.1% formic acid and loaded onto a self-packed C18 StageTip^63^ (five discs, Sigma Aldrich). Tips were equilibrated with 0.1% formic acid. After loading, tips were washed four times with 200 µL 50 mM ammonium formate (pH 10) in water. Peptides were eluted with 150 µL 50 mM ammonium formate (pH 10) in 35% acetonitrile. For all other samples, volumes were adjusted to 150 µL (3-dish) or 600 µL (12-dish) with acetonitrile and formic acid to final concentrations of 30% acetonitrile and 0.1% formic acid, and samples were centrifuged at 20000 g for 5 min at room temperature. Ninety percent of the volume of each sample was transferred to a fresh tube and loaded onto a self-packed SCX StageTip (two discs, Sigma Aldrich). Tips were equilibrated with 0.1% formic acid in 30% acetonitrile. After loading, tips were washed four times with 200 µL 0.1% formic acid in 30% acetonitrile. Peptides were eluted with 150 µL 100 mM ammonium formate (pH 6.5) in 50% acetonitrile. Finally, all StageTip eluates were lyophilized in a SpeedVac for LC-MS/MS analysis.

### LC-MS/MS analysis

LC-MS/MS samples were measured using one of three LC-MS settings. Setting A: an Orbitrap Fusion Lumos Tribrid mass spectrometer (Thermo Fisher Scientific) connected to a Dionex UltiMate 3000 RSLCnano system (Thermo Fisher Scientific) equipped with an in-house packed trap column (2 cm x 75 μm) packed with 5 μm C18 resin (Reprosil PUR AQ - Dr. Maisch) and an analytical column (48 cm x 75 μm, heated to 55°C) packed with 3 μm C18 resin (Reprosil PUR AQ - Dr. Maisch). Solvent A consisted of 0.1% formic acid and 5% DMSO in water; solvent B consisted of 0.1% formic acid and 5% DMSO in acetonitrile; solvent C consisted of 0.1% formic acid in water. Orbitrap resolution values are reported relative to m/z 200. Setting A was used for all samples in Figure 1 and 2, and for tuning the RNA length with different nucleases (Figure 5A and S5A). Setting B: an Orbitrap Astral mass spectrometer (Thermo Fisher Scientific) connected to an Evosep One (Evosep) chromatography system equipped with an Aurora Rapid 8×75 XT C18 UHPLC column (8 cm x 75 μm, 1.7 μm C18, heated to 50°C). Solvent A consisted of 0.1% formic acid in water; solvent B consisted of 0.1% formic acid in acetonitrile. Setting B was used for crosslinking site localization samples (Figure 3 and 4). Setting C: an Orbitrap Eclipse Tribrid mass spectrometer (Thermo Fisher Scientific) with the same LC system as in Setting A. Setting C was used for dual sequencing samples (Figure 5 and 6).

#### LC-MS/MS for RNA digest spike-in experiments and peptide-RNA crosslinks

For LC-MS setting A, lyophilized HeLa peptide standard with or without RNA digest spike-in and peptide-RNA crosslink samples were reconstituted in 12 µL solvent C and 10 μL were injected per run. After injection, the sample was transferred onto the trap column and then washed with C at 5 μL/min for 10 min before transfer to the analytical column using a gradient of solvent A and solvent B. Separation was performed on an 80 min (12-dish peptide-RNA crosslink sample for ID comparison in Figure 2F-2I and Figure S2C) or a 50 min (HeLa peptide standard and other peptide-RNA crosslink samples) linear gradient with a flow rate of 300 nL/min from 4% solvent B to 32% solvent B. The nanospray source voltage was 2100 V, and the ion transfer tube temperature was 275°C. Detection occurred with data-dependent acquisition using an OTOT method and a cycle time of 2 s. Full MS scans (MS1) were acquired at a resolution of 60000, scan range 360-1300, RF lens 50%, normalized AGC target 100% and maximum injection time 50 ms. Based on the MS1s, precursors were targeted for MS2 scans if the charge was between 2 and 6, the monoisotopic peak determination (MIPS) was peptide, and the intensity exceeded 2.5E4. MS2 isolation occurred with a quadrupole isolation window of 1.7 Th and fragmentation with 30% HCD collision energy. MS2 spectra were acquired at a resolution of 15000 (HeLa peptide standard) or 30000 (peptide-RNA crosslink samples), first mass 100 Th, normalized AGC target 300% (HeLa peptide standard) or 100% (peptide-RNA crosslink samples), and maximum injection time 25 ms (HeLa peptide standard) or 50 ms (peptide-RNA crosslink samples). Dynamic exclusion was enabled with an exclusion duration of 20 s following precursor fragmentation.

For LC-MS setting B, lyophilized peptide-RNA crosslink samples were reconstituted in 100 µL 0.1% FA and loaded onto Evotips (Evosep) according to the manufacturer’s instructions. The samples were measured using the Whisper Zoom 20 samples-per-day (SPD) method with an MS acquisition time of 68.6 minutes. The nanospray source voltage was 2000 V, and the ion transfer tube temperature was 275°C. Detection occurred with data-dependent acquisition using an OTAT method and a cycle time of 0.6 s. Full MS scans (MS1) were acquired at a resolution of 240000, scan range 360-1300, RF lens 50%, normalized AGC target 300% and maximum injection time 10 ms. Based on the MS1s, precursors were targeted for MS2 scans if the charge was between 2 and 6, and the monoisotopic peak determination (MIPS) was peptide. MS2 isolation occurred with a quadrupole isolation window of 1.2 Th and fragmentation with 30% HCD collision energy. MS2 spectra were acquired at a scan range of 100-2000, normalized AGC target 300%, and maximum injection time 3.5 ms. Dynamic exclusion was enabled with an exclusion duration of 45 s following precursor fragmentation.

#### Collision energy ramp for oligonucleotides and peptides

Twenty-four ng RNA oligo (5’-UAGUAG-3’, dissolved in 10 µL 0.1% formic acid), 100 fmol PROCAL^44^ (dissolved in 1 µL 0.1% formic acid) or 50 ng HeLa peptide standard (dissolved in 1 µL 0.1% formic acid) were injected per run. After injection, the sample was transferred onto the trap column and then washed with C at 5 µL/min for 5 min before transfer to the analytical column using a gradient of solvent A and solvent B. Separation was performed on a 9 min (RNA oligo) linear gradient from 4% solvent B to 20% solvent B or a 22 min gradient (PROCAL and HeLa peptide standard) with a flow rate of 300 nL/min. The nanospray source voltage was 2100 V, and the ion transfer tube temperature was 275°C. Detection occurred with data-dependent acquisition using an OTOT method and a cycle time of 2 s.

For the RNA oligo, MS1 acquisition used quadrupole-isolated targeted selected ion monitoring (tSIM) over the m/z range 1000-2000 Th at a resolution of 60000, RF lens 100%, normalized AGC target 200% and maximum injection time 50 ms. Based on the MS1s, precursors were targeted for MS2 scans if the charge was between 1 and 6. MS2 isolation occurred with a quadrupole isolation window of 1.3 Th and HCD collision energy was stepped from 10% to 30% in 2% increments. MS2 spectra were acquired at a resolution of 30000, first mass 100 Th, normalized AGC target 300%, and maximum injection time 100 ms. Dynamic exclusion was enabled with an exclusion duration of 10 s following precursor fragmentation.

For PROCAL and the HeLa peptide standard, full MS scans (MS1) were acquired at a resolution of 60000, scan range 360-1300, RF lens 50%, normalized AGC target 100% and maximum injection time 50 ms. Based on the MS1s, precursors were targeted for MS2 scans if the charge was between 2 and 6, the MIPS was peptide, and the intensity exceeded 2.5E4. MS2 isolation occurred with a quadrupole isolation window of 1.7 Th and HCD collision energy was stepped from 10% to 30% in 2% increments. MS2 spectra were acquired at a resolution of 15000, first mass 100 Th, normalized AGC target 300%, and maximum injection time 25 ms. Dynamic exclusion was enabled with an exclusion duration of 20 s following precursor fragmentation.

#### Dual-MS2 method for sequential sequencing of peptide-RNA crosslinks

For LC-MS setting C, lyophilized peptide-RNA crosslinks produced with benzonase-digestion from three 150 mm dishes of MCF7 cells were reconstituted in 12 µL solvent C and 10 µL were injected. After injection, the sample was transferred onto the trap column and then washed with C at 5 µL/min for 10 min before transfer to the analytical column using a gradient of solvent A and solvent B. Separation was performed on a 50 min linear gradient with a flow rate of 300 nL/min from 4% solvent B to 32% solvent B. The nanospray source voltage was 2100 V, and the ion transfer tube temperature was 275°C. Detection occurred with data-dependent acquisition using an OTOT method and a cycle time of 2 s.

Full MS scans (MS1) were acquired at a resolution of 60000, scan range 350-2000, RF lens 40%, normalized AGC target 100% and maximum injection time 50 ms. Based on the MS1s, each precursor was targeted for two dependent MS2 scans with a quadrupole isolation window of 1 Th if the charge was between 2 and 6, the MIPS was peptide, and the intensity exceeded 2.5E4. MS2s for peptide sequencing were acquired with activation type HCD, collision energy 30% and first mass 100 Th. MS2s for RNA sequencing were acquired with activation type CID, collision energy 22%, CID activation time 30 ms, activation Q 0.15 and scan range 200-2000 Th. All MS2 spectra were acquired at a resolution of 30000, normalized AGC target 200%, and maximum injection time 54 ms. Dynamic exclusion was enabled with an exclusion duration of 20 s following precursor fragmentation.

### Data analysis

Proteomic data were processed in R (version 4.3.0) within RStudio (version 2024.12.0) or in Python (version 3.14 by default, 3.12 for FINCHES).

#### Database search for peptide identification

All data were searched against the human UniProt database^64^ (SwissProt reviewed, from December 2022, 20,434 entries). MaxQuant^65^ (version 2.2.0.0) was used for searching HeLa peptide standards with RNA spike-ins, using default values for all parameters.

All other data were searched with MSFragger^66^ (version 4.3) within FragPipe (version 23.1) for peptide-RNA crosslink identification and quantification, together with IonQuant^67^ (version 1.11.11) and Philosopher^68^ (version 5.1.2). MS2 spectra were searched against the human UniProt database supplemented with common contaminants and a 50% decoy set using the built-in ‘Add decoys’ function in FragPipe. Default parameters for the built-in workflow ‘XRNAX-MassOffset’ were applied unless stated otherwise. Enzymatic specificity was set to ‘stricttrypsin’ with up to 2 missed cleavages allowed. Peptide length was set to 2 to 50 amino acids with a peptide mass range of 50-5000 Da. Mass tolerances were set to 10 ppm for precursor and 20 ppm for fragment, with isotope error correction of 0/1/2. For all pepR-MS samples, no fixed modification was set and variable modifications included oxidation (M) and protein N-terminal acetylation. For searches of representative MS data from previous studies, carbamidomethyl (C) was set as a fixed modification, or no modification was set depending on their workflow (for details see parameter files deposited to MassIVE); variable modifications were set to oxidation (M) and protein N-terminal acetylation. The maximum number of variable modifications per peptide was set to 3. Mass offset searching was enabled with modifications allowed at all amino acids and a mass offset list customized for each workflow (pepR-MS or from previous studies). In general, for all pepR-MS MS data, mass offsets were generated by searching representative files with all possible mass offsets up to one nucleotide and the rarely identified mass offsets were filtered out. Based on the most frequently identified mass offsets up to one nucleotide (0, 94.0167, 288.0147, 306.0253, 322.0025, and 340.013we generated all possible masses of RNA chains up to three (default if not specified otherwise) or seven (for the tuneable length with nucleases and dual sequencing samples, Figure 5 and 6) nucleotides. For the crosslinking site localization samples (Figure 3 and 4), only 0 and 2-nt mass offsets were used, as nuclease P1 digestion predominantly produces 2-nt RNA moieties. For representative MS data from previous studies, a similar procedure was performed to generate mass offsets with up to three nucleotides for each raw file from Kramer *et al.*, 2014^7^ and Trendel *et al.*, 2019^10^, only considering uridine as the crosslinking base. For Bae *et al.*^16,17^ mass offsets were directly adapted from the search parameters used in the original study. All mass offsets can be found in the search parameter files deposited to MassIVE. Labile-mode searching^69^ was enabled for all mass offsets with remainder fragment masses of one nucleotide (default) or mass offsets for a single base (for HF-digested files). PSM validation used PeptideProphet and protein inference used ProteinProphet with default parameters for offset search. Quantification was performed with IonQuant using LFQ (MaxLFQ calculated with min. 2 ions) and match-between-runs set to off.

#### Definition of crosslink identifications

For all samples except for the dual sequencing ones, the count of identified crosslinks was defined as the number of unique modified peptides crosslinked to unique RNA mass offsets at specific amino acid positions in the peptide sequence (e.g. peptide PEPTIDEK crosslinked to an RNA of mass 306 at position P3). For the dual sequencing samples, the crosslink count was defined as the number of unique modified peptide sequences crosslinked to unique RNA sequences (e.g. peptide PEPTIDEK crosslinked to RNA XUG). The count of identified crosslinking sites was defined as the number of unique crosslinking amino acid positions within the full-length protein sequence, regardless of the crosslinked RNA (e.g. G1158 in the protein DHX9). Crosslinking site ID in Figure 2 was based on reported modification sites by MSFragger as Assigned Modifications, whereas the (focused) crosslinking sites in Figure 3 and 4 were based on aggregated PSM evidence as described below.

#### Crosslinking site analysis

Crosslinking site analysis (Figure 3 and 4) was based on three biological replicates, each separated into nuclear and cytosolic fractions. PSMs from the same protein were aggregated for better crosslinking site localization. Every PSM bearing an RNA moiety was considered one unit of evidence and was assigned candidate crosslink positions reported by MSFragger. Because this assignment was frequently ambiguous, the PSM evidence unit was distributed evenly across all its candidate residues (a PSM with n candidate sites therefore contributed 1/n PSM-units to each residue, or one PSM-unit to a unique candidate). Precursor intensities were distributed in parallel using the same 1/n split for the intensity analysis in Figure 3G. PSM-units were then summed over PSMs of the same peptide in all replicates, and finally over all peptides of the same protein. This yielded, for every protein, a single-residue-resolution profile of PSM-units along the protein sequence. Residues were first clustered along the sequence, bridging gaps of up to two unmodified residues and splitting the profile wherever a larger gap occurred; each cluster was zero-padded by two positions on either side so that terminal residues remained eligible for local maximum detection. Local maxima with at least one PSM-unit were then detected using SciPy’s (version 1.17.1) scipy.signal.find_peaks; clusters containing no qualifying maximum were discarded as noise. Where a cluster contained several local maxima, boundaries were placed at the minimum of the profile between consecutive peaks, and each residue was assigned to the peak on its side of that boundary. Each local maximum defined one crosslinking site, reported with its residue composition, summed PSM-units, number of contributing peptides, and the identity of the residue(s). When two or more residues shared the identical maximum PSM-unit value, a single representative position was assigned per site as the median of the tied positions. Position-based annotations were performed on this representative position, each site thus contributing exactly one crosslinking position. Analyses that depend on the chemical identity of the crosslinked residue were restricted to uniquely localized sites, defined as those with exactly one peak residue.

Statistical enrichment or depletion of amino acids at crosslinking sites (Figure 3B) was tested using two-sided Fisher’s exact tests based on site counts, comparing amino acid frequencies at crosslinking sites to their overall frequencies in the crosslinked protein sequences. P-values were corrected for multiple testing using the Benjamini-Hochberg procedure.

Gene ontology (GO) term analysis was performed using the DAVID^70^ (version 6.8) functional annotation tool, using a deep MCF7 full proteome as background^71^.

#### Protein domain annotation and RRM analysis

Domain annotations for crosslinked proteins were obtained from InterPro^72^ (version 92.0 to match the protein sequence database from December 2022). The list of human ribosomal proteins was downloaded from UniProt (189 entries, January 2025). To generate a viable AlphaFold^20^ prediction of a generic RRM domain (Figure 4F), the Pfam consensus sequence needed to be extended on both sides. Only the consensus sequence extracted from the Pfam profile hidden Markov model (HMM) appeared truncated and lacked some secondary structure elements upon AlphaFold^20^ prediction. To extend the profile HMM, the seed alignment of 68 sequences of the Pfam entry PF00076 was downloaded. These sequences were mapped back to their original protein sequences and extended at both the N and C-termini by five amino acids. The extended sequences were aligned to the original profile HMM using the hmmalign function in HMMER^73^ (version 3.3) and the new alignment was used to build an extended HMM using hmmbuild. Crosslink annotations for RRMs were refined by searching all crosslinked proteins with the extended HMM using hmmsearch. Sequences of RRM domains annotated in this step were realigned with hmmalign, and crosslinks were mapped to this updated alignment for Figure 4E and 4F. The extended HMM was then used to generate sequence logos for the RRM, and its consensus sequence alongside the RNA sequence 5’-CAUC-3’ was provided as input for AlphaFold 3^20^ structure prediction. Positions of secondary structures from the predicted RRM consensus (Figures 4E and S4D) were extracted using DSSP 4^74^.

#### IDR annotation and local sequence context analysis

Disordered regions were predicted with MobiDB-lite^75^ (version 4.0) from the protein sequences in the search database. Only IDRs of at least 30 consecutive residues were used for analysis. Crosslinking sites were grouped into four classes based on their positions. Sites inside any InterPro domain were classified as in-domain. Of the remainder, sites inside a predicted IDR were classified as in-IDR if the site lay ≥10 aa from the nearest InterPro domain, and domain-close if it lay <10 aa from the nearest domain edge; the distance was measured from the edges of PSM evidence distribution (local minima) to the nearest domain edge on the same protein, and was 0 for overlap. All other sites were classified as no-annotation. Domain-close and no-annotation sites were excluded from the analyses, except for intensity analysis in Figure 3G, where domain-close sites were included in the IDR group.

IDRs (≥30 aa) and RNA-binding domains were classified as crosslinked if at least one crosslinking site assigned to that region class (in-IDR or in-domain) fell within the region, and non-crosslinked otherwise. Within each region class, residues of crosslinked proteins inside a region were divided into crosslinked residues (or crosslinking sites, i.e. local maxima of aggregated PSM evidence), non-crosslinked residues in crosslinked regions, and non-crosslinked residues in non-crosslinked regions.

Crosslink spacing was analysed separately within IDRs and RBDs containing at least two crosslinking sites. For each crosslinking site, the nearest-neighbour distance was calculated as the smallest distance to another site within the same region. Background distributions were generated using 10,000 random configurations per region, preserving crosslinking site number and crosslinked amino acid composition. Positions were sampled only from theoretically trypsin-detectable residues. Detectability was defined using an in-silico digest of the full protein sequence, with cleavage after lysine or arginine except before proline, allowing up to two missed cleavages and peptide lengths of 7-47 amino acids. Clustering was assessed using a region-specific rank statistic, defined as the probability that an observed nearest-neighbour distance was shorter than a randomized distance from the same region (AUC > 0.5 indicates shorter-than-expected spacing, i.e. clustering). These AUC values were averaged across regions. The null distribution was constructed by similarly averaging randomized configuration AUCs. Significance was assessed using the two-sided empirical permutation p-value with the +1 correction (*p* = (2 · min(tail) + 1)/(*N* + 1)), where *N* is the number of permutations). P-values were floored at 9.99E-5 with 10,000 permutations. IDRs and RBDs were tested separately without multiple-testing correction. For visualization, observed and randomized distances were pooled separately within each region type. The brackets report the two global equal-region AUC permutation tests described above (***: p < 1E-4).

NARDINI+/GIN clusters were assigned as described for the NARDINI+_from_FASTA notebook^32^ (version 1.2), reimplemented for batch processing, using MobiDB-lite-predicted IDRs with the *Homo sapiens* IDRome from the original publication as the compositional prior, and 10,000 sequence scrambles per IDR as the patterning null. For each of the 30 GIN clusters, the enrichment of crosslinked IDRs against non-crosslinked IDRs was tested with a two-sided Fisher’s exact test, and the enrichment reported as a log2 odds ratio with a 0.5 pseudocount. The 30 p-values were corrected across clusters by the Benjamini-Hochberg procedure.

Sequence composition around crosslinking sites was analysed separately for IDRs and structured domains. A crosslink window spanned five residues on either side of a crosslinking site. Windows extending beyond a protein terminus were excluded. Background windows comprised every 11-AA window of a region, taken at single-residue steps. Because the crosslinked residue would otherwise contribute to the window composition, the centre residue was removed from every window before calculating composition, so each window contributed its ten flanking residues, and the comparison reflected sequence context only. To remove crosslink-associated sequence from the background, every sliding window overlapping any assigned crosslink’s full 11-residue span was discarded. For regions containing multiple crosslinks, this exclusion was applied to all their crosslink spans. A second background comprised sliding windows from non-crosslinked regions. For each window, the fraction of each of the 20 amino acids was calculated after centre removal. The heatmaps in Figure 4C compared crosslink-centred windows with crosslinked-region background. Heatmap colour represented log₂ fold change. Cell labels reported Benjamini-Hochberg-adjusted q-values from two-sided Mann-Whitney U tests comparing the per-window fractions. Correction was applied across the 20 amino-acid tests separately for the IDR and domain analyses. In Figure S4C, three distributions were compared: crosslink-centred windows, windows from crosslink-excised crosslinked regions and windows from non-crosslinked regions. Each window contributed counts among its ten retained residues for five amino-acid groups: aromatic (FHWY), negatively charged (DE), positively charged (KR), polar (CNQST) and aliphatic (AGILMPV).

Sequence logos around crosslinking sites were generated separately for IDRs and structured domains. A crosslink window spanned ten residues on either side of a crosslinking site. Crosslinking sites were kept only if they were uniquely localized and were discarded if the window would extend beyond a protein terminus. For each class (IDRs and structured domains), a background residue composition was pooled over the corresponding regions of all crosslinked proteins. These pools gave one position-independent amino-acid (or group) frequency vector per class. Amino-acid information-content logos were generated using Logomaker’s (version 0.8.7) alignment_to_matrix, transform_matrix and Logo functions. Reference compositions were calculated by pooling residues across all annotated IDRs or domains of all crosslinked proteins. For grouped enrichment plots, residues were assigned to the amino acid groups defined above. At each aligned position, each group’s frequency was compared with the corresponding pooled IDR or domain background using a log₂ odds ratio with a 0.5 pseudocount. Values were capped at ±4 and displayed as grouped bars.

IDR sequences were annotated with ten RNA-binding motifs: RG/RGG boxes, [R/K]S repeats, AT-hooks, [F/Y]GG boxes, K patches, R patches, G patches, S patches, Q/N patches and aromatic patches. The definitions were designed based on real sequences and/or adapted from the NARDINI+^32^ algorithm:

- RG/RGG box: At least two RG, RGG or RGGG units separated by up to four non-arginine residues.

- [R/K]S repeat: At least three consecutive [R/K]S or S[R/K] dipeptides.

- AT-hook: A GRP core with K/R within three residues on either side.

- [F/Y]GG box: Individual [F/Y]GG triplets and repeat boxes whose start positions were no more than ten residues apart.

- R, K, G and S patches: At least four occurrences of the specified residue, separated by no more than two other residues.

- Q/N and aromatic patches: The same patch rule applied jointly to Q/N or F/Y/H/W, respectively.

Short linear motifs (SLiMs) were predicted by submitting IDR sequences to the Eukaryotic Linear Motif (ELM) sequence-search API^76^. Sequences longer than 1,500 residues were divided into windows with ten-residue overlaps. All returned predictions were retained; experimentally validated annotation was not required. The figure included seven representative ELM classes: MOD_CK1_1, MOD_CK2_1, MOD_GSK3_1, MOD_PKA_1, MOD_PKA_2, MOD_SUMO_for_1 and MOD_SUMO_rev_2. For each residue, distance to a motif was defined as the minimum distance to the nearest motif boundary within the same IDR. Distance was zero inside a motif. IDRs lacking the corresponding motif were excluded from that motif’s analysis. Non-crosslinked IDRs were not used in the comparison. For each motif, distances from crosslinked residues were compared with distances from all non-crosslinked residues in the same IDRs. The difference was summarized using a Mann-Whitney-based AUC statistic, which measures how often a crosslinked residue is closer to the motif than a background residue. Significance was assessed using 10,000 randomizations. Within each IDR, the observed number of crosslinked residues was preserved, but their positions were randomly reassigned. The randomly selected residues were compared with the remaining background residues, and the AUC was recalculated. The observed AUC was then compared with the randomized AUC distribution to obtain a two-sided empirical p-value. Benjamini-Hochberg correction was applied across the 17 selected motifs (*: p < 0.05).

Interactions between 101 crosslinked IDRs and 1,739 IDRs from MCF7 RNA-binding proteins were predicted using FINCHES^40^ (version 0.1.2), specifically the Mpipi_frontend.intermolecular_idr_matrix function, with an 11-AA window and default model settings. Partner IDRs were MobiDB-lite regions of at least 30 residues from proteins annotated with GO:0003723 (“RNA binding”). Attractive and repulsive profiles were calculated using the FINCHES vector algorithms applied to predicted interaction matrices, with a zero threshold and mean aggregation. Profiles were smoothed using a five-position, second-order Savitzky-Golay filter and averaged equally across partner IDRs. For the YBX1 site map, consecutive positions with mean attractive ε < −1 or mean repulsive ε > +1 were labelled as candidate sticker or spacer regions, respectively. The accompanying heatmap showed interactions averaged over each partner’s windows, with partners ordered by Euclidean distance/Ward hierarchical clustering. For the distribution plots, scores were pooled across all evaluated positions and displayed alongside the crosslinked subset using SciPy’s gaussian_kde with bandwidth factor 0.25. Crosslinked and non-crosslinked residues of crosslinked IDRs were compared using two-sided Mann-Whitney U tests.

#### RNA identification in dual sequencing

Dual sequencing samples were measured with the sequential MS2 method and peptides were first identified from peptide-sequencing MS2 spectra (HCD) using MSFragger as described above. The paired RNA-sequencing MS2 spectra (low-q CID) were analysed using SALT v0.1.0. To recover isotope envelopes removed from the MSFragger-calibrated, deisotoped mzML files, Thermo RAW files were reconverted with ProteoWizard msconvert (version 3.0.24040-1c422c5) using vendor peak picking. Peaks in the resulting uncalibrated mzML spectra were matched scan-wise to calibrated peaks by identical intensity, and the corresponding mass corrections were transferred to the complete raw peak list. RNA-sequencing MS2 spectra were then charge-deconvoluted with the pyOpenMS^77^ Deisotoper (version 3.5.0) using a 10-ppm isotope tolerance, charge states 1-6, 3-10 isotope peaks, the decreasing-intensity model, summed isotope-envelope intensities, and conversion to singly charged peaks. Every PSM of crosslinked peptides was paired by precursor m/z with its corresponding RNA-sequencing spectrum. The unmodified peptide mass was calculated as the MSFragger-calculated precursor mass minus the assigned delta mass and subtracted from the deconvoluted RNA-sequencing spectrum to obtain RNA-fragment ladders. Each spectrum was further calibrated against the peak within ±0.05 Da of the 95.024538-Da uracil-derived marker ion, with the resulting mass shift applied to all peaks. The theoretical library comprised every 2-5-nt sequence containing one designated crosslink position, X, with the remaining positions drawn from A, U, G, C and S (4SU). Single phosphodiester-bond cleavages generated peptide-retaining (primary) and non-peptide-retaining diagnostic fragments, whereas multiple cleavages generated secondary peptide-retaining fragments. For false-discovery assessment, noncanonical decoy fragments were generated by adding 41.001397 Da (azido) per nucleotide; the analysis used the combined target-azido library containing equal numbers of target and decoy sequences. Candidate RNA sequences were first restricted to the RNA length inferred from the precursor offset mass, and observed peaks were matched to theoretical primary, secondary and diagnostic fragments within 0.01 Da. When several observed peaks matched one theoretical fragment, the most intense was selected. Candidates containing at least one matched crosslink-retaining fragment were scored separately for the three fragment classes *H* = ln(*n*! ∑ *I_rel_*), where *n* is the number of matched fragments and *I_rel_* is relative intensity. Candidates were ranked first by primary Hyperscores. Ties were resolved sequentially by secondary and diagnostic Hyperscores, with unresolved ties retained as ambiguous assignments. The delta Hyperscore was the difference between the best and next distinct lower primary Hyperscore. Identification confidence was controlled separately for each experiment using target-decoy competition. Scan-level target and decoy winners were ranked by decreasing primary Hyperscore. At each score threshold, the false discovery rate (FDR) among target identifications was estimated as (*D* + 1)/*T*, where *T* and *D* are the cumulative numbers of target and decoy winners, respectively^78,79^. Threshold-specific estimates were converted to monotonic q-values, and only target identifications with *q* ≤ 0.01 were retained and joined to their corresponding PSMs to report both peptide and RNA sequences. Per-scan E-values were subsequently calculated for FDR-surviving identifications according to Fenyö and Beavis^80^ . The tail threshold for the empirical survival function was increased from 0.1 in the original method to 0.2 because the number of RNA sequence candidates per scan was smaller than in peptide-sequence searches, and a threshold of 0.1 generally provided too few tail points for a stable fit.

#### Data visualization

Schematics in Figure 1, 2, and S2 were partially created with BioRender. Data were plotted using the R package ggplot2^81^ (version 3.5.2) and the Python package Matplotlib^82^ (version 3.10.8). Protein structures were visualized in UCSF ChimeraX^83^ (version 1.7.1). Chemical structures were drawn with ChemDraw (version 21.0.0). Figures were composed in Adobe Illustrator 2024.

## Data availability

Proteomic data and search results have been deposited in the MassIVE database under the identifier MSV000097992. SALT is available on GitHub (https://github.com/kusterlab/SALT).

## Acknowledgements

We thank all members of the Kuster laboratory for continuous support and discussion. We gratefully acknowledge funding by the German Research Council (DFG) supporting this work (project number 325871075 (SFB1309-B02) and project number 492625837).

## Author contributions

Conceptualization, J.T., B.K.; methodology, S.S., J.T.; investigation, S.S.; writing—original draft, S.S., J.T., B.K.; resources, B.K.; data curation, S.S., J.T.; writing—review and editing, S.S., J.T., B.K.; visualization, S.S.; supervision, J.T., B.K.; project administration, J.T.; funding acquisition, J.T., B.K..

---

**Figure S1:**
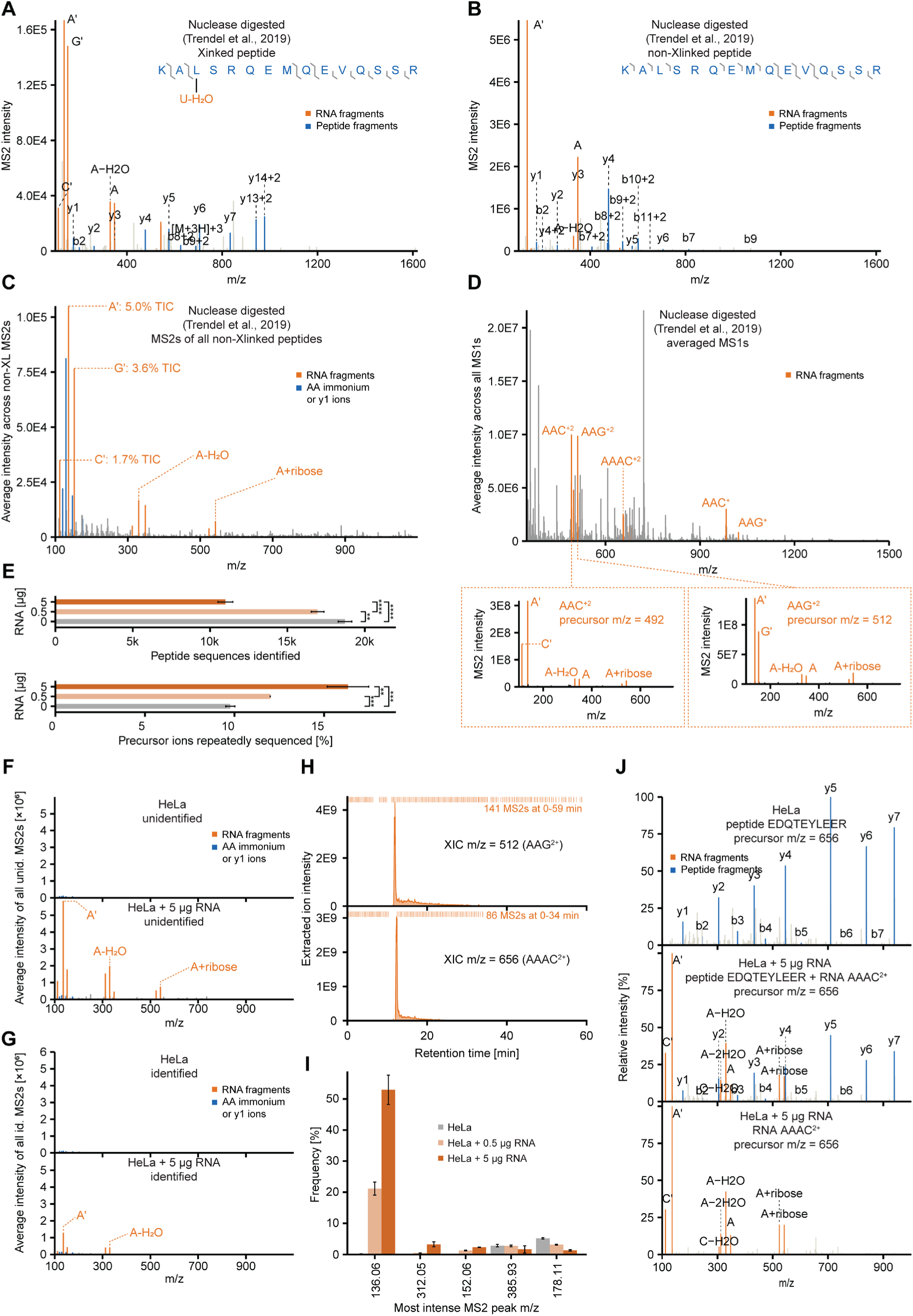
Non-crosslinked RNA contamination in isolates of peptide-RNA crosslinks. **A-D.** Published LC-MS data of nucleotide-crosslinked peptides using a nuclease-based isolation procedure (Trendel *et al.*, 2019). **A.** Exemplary MS2 spectrum of a peptide-RNA crosslink showing contamination by foreign RNA fragments. **B.** Same as in panel A but for peptides identified as unmodified (non-crosslinked). **C.** Averaged MS2 spectra of all peptides identified as unmodified (non-crosslinked). **D.** Averaged MS1 spectra of the entire MS run. Inserts show corresponding MS2 spectra of two highlighted RNA fragments. **E-I.** LC-MS data from this study using a HeLa standard with or without nuclease-digested RNA spike-ins. **E.** Bar graph showing number of identified peptide sequences (top) and the proportion of repeatedly sequenced MS1 peaks (bottom), related to Figure 1C-E. Statistical analysis was performed using a two-tailed unpaired Student’s t-test. **, p < 0.01; ***, p < 0.001; ****, p < 0.0001. Error bars indicate standard deviation across triplicates. **F.** Average of identified MS2 spectra for each LC-MS run. **G.** Same as in panel F but for unidentified MS2 spectra. **H.** Extracted ion chromatograms (XIC) of exemplary RNA-derived precursors. MS2 spectra of those precursors are annotated. **I.** Bar graph comparing the occurrence of the most intense MS2 peaks in one MS run. Percentages indicate the fraction of MS2 spectra, among all acquired MS2 spectra, in which a peak at the specified mass was the most intense peak (base peak). Error bars indicate standard deviation across triplicates. **J.** Exemplary MS2 spectra of precursors with the same m/z and retention time in LC-MS runs with and without RNA spike-in.

**Figure S2:**
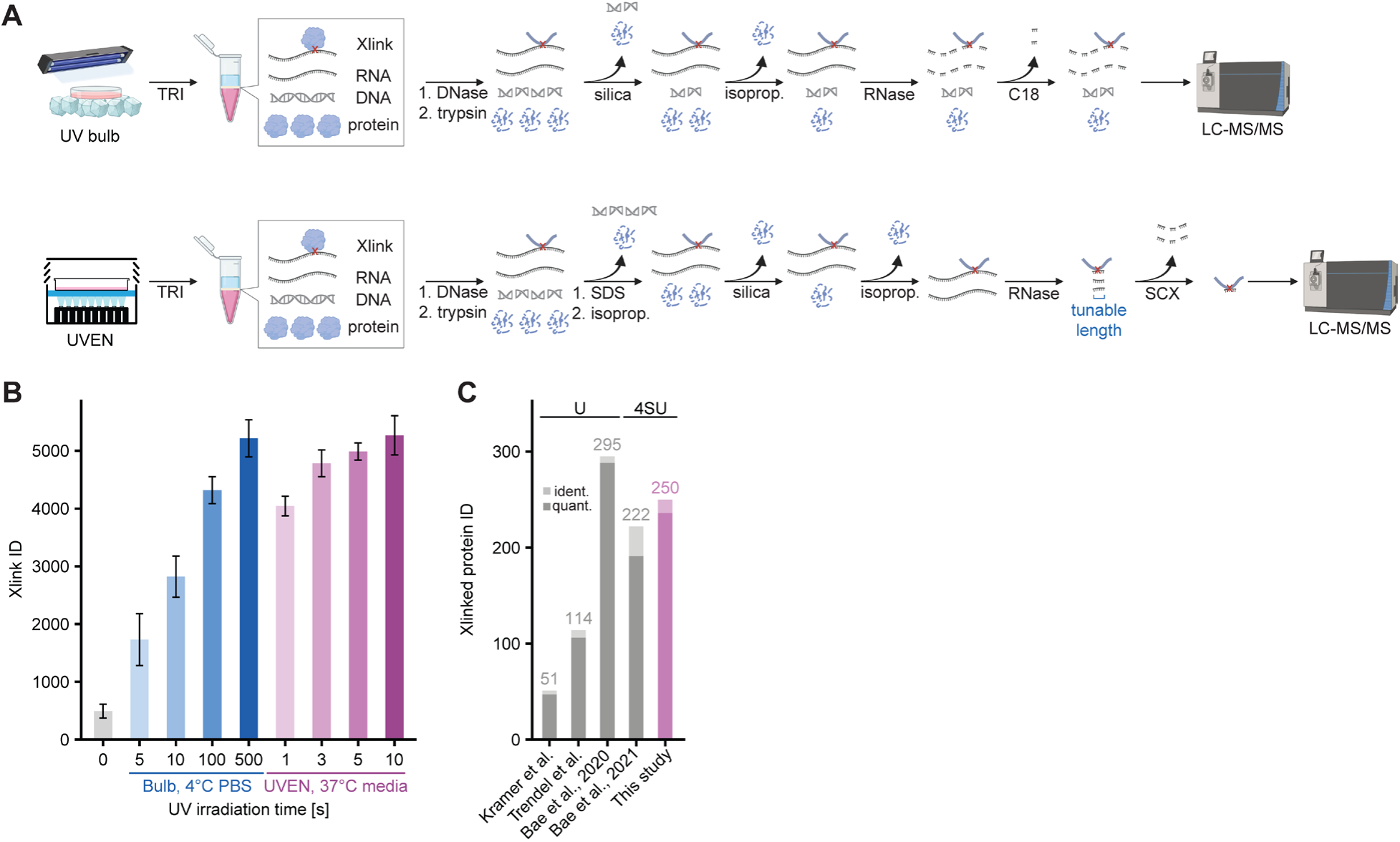
PepR-MS identifies peptide-RNA crosslinks with enhanced robustness and sensitivity. **A.** Detailed experimental scheme for the purification of peptide-nucleotide crosslinks with (lower panel) and without (upper panel) dedicated removal of non-crosslinked RNA. **B.** Bar graph showing identified crosslinks using the high-power UV irradiation device (UVEN) and conventional UV bulb (Vilber crosslinker). Error bars indicate standard deviation across triplicates.**C.** Identified and quantified crosslinked proteins in previous studies and this study, using 365 nm crosslinking (main crosslinking base 4SU) or 254 nm crosslinking (main crosslinking base U).

**Figure S3.**
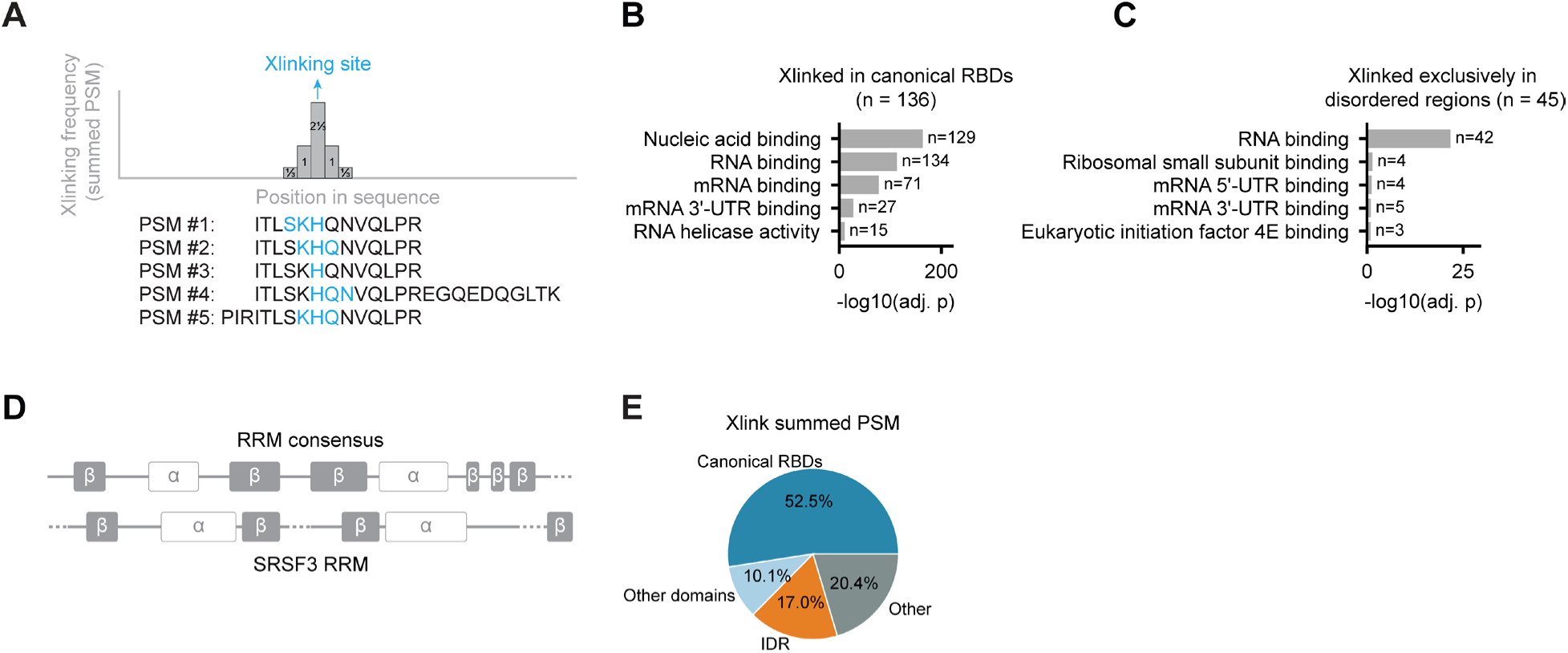
RNA crosslinking sites reveal residue and domain level preferences. **A.** Schematic of the crosslinking site refinement method. **B-C.** Top five enriched Gene Ontology (GO) terms of (B) proteins crosslinked within canonical RNA-binding domains (same as in Figure 3G) and of (C) proteins that crosslinked exclusively through disordered regions predicted by MobiDB-lite. **D.** Alignment of RRM secondary structures in panel F (top: Pfam domain definition; bottom: Reference NMR solution structure of SRSF3 RRM (PDB 2I2Y)) **E.** Pie charts showing the fraction of summed PSMs contributed by crosslinks in canonical RNA-binding domains (same as in S3B), other InterPro domains, IDRs and other regions.

**Figure S4.**
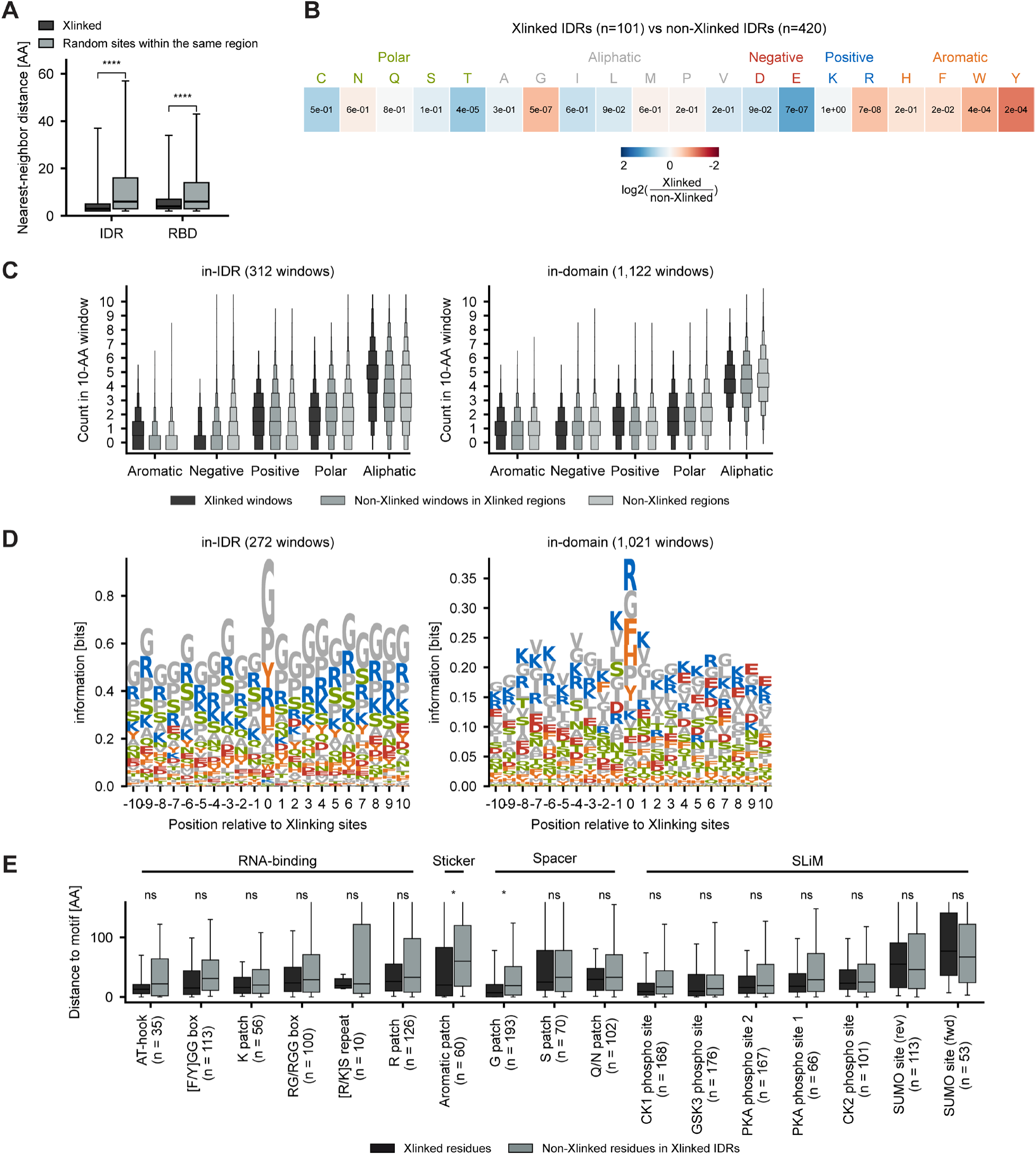
RNA crosslinking sites show IDR-specific local sequence features absent from structured domains. **A.** Box plots comparing nearest-neighbour distances between crosslinking sites with those between amino-acid-matched sites randomly drawn from the same IDRs (left) and RNA-binding domains (right). Significance was assessed using permutation tests within the same region. ****, p < 1E-4. **B.** Heatmap of the difference in amino-acid composition of crosslinked and non-crosslinked IDRs of all crosslinked proteins. Colour indicates log2 fold-change in AA fraction; cells are annotated with p-values calculated using a two-sided Mann-Whitney U test and adjusted using the Benjamini-Hochberg method. **C.** Histogram-violin plots comparing AA-group composition across three sets of 10-AA windows in IDRs (left) and domains (right): windows centred on crosslinking sites (Xlinked windows), on non-crosslinked positions in crosslinked IDRs, and on non-crosslinked positions in non-crosslinked IDRs. **D.** Sequence logo of amino-acid composition across the ±10-AA window centred on crosslinking sites in IDRs (left) and domains (right). **E.** Box plots comparing distances from crosslinking sites to the nearest annotated sequence feature with those from non-crosslinked residues in the same crosslink-containing IDRs, across RBP-associated motifs, sticker and spacer patches, and short linear motifs. Significance was assessed using within-IDR permutation tests with Benjamini-Hochberg correction. *, adjusted p < 0.05.

**Figure S5:**
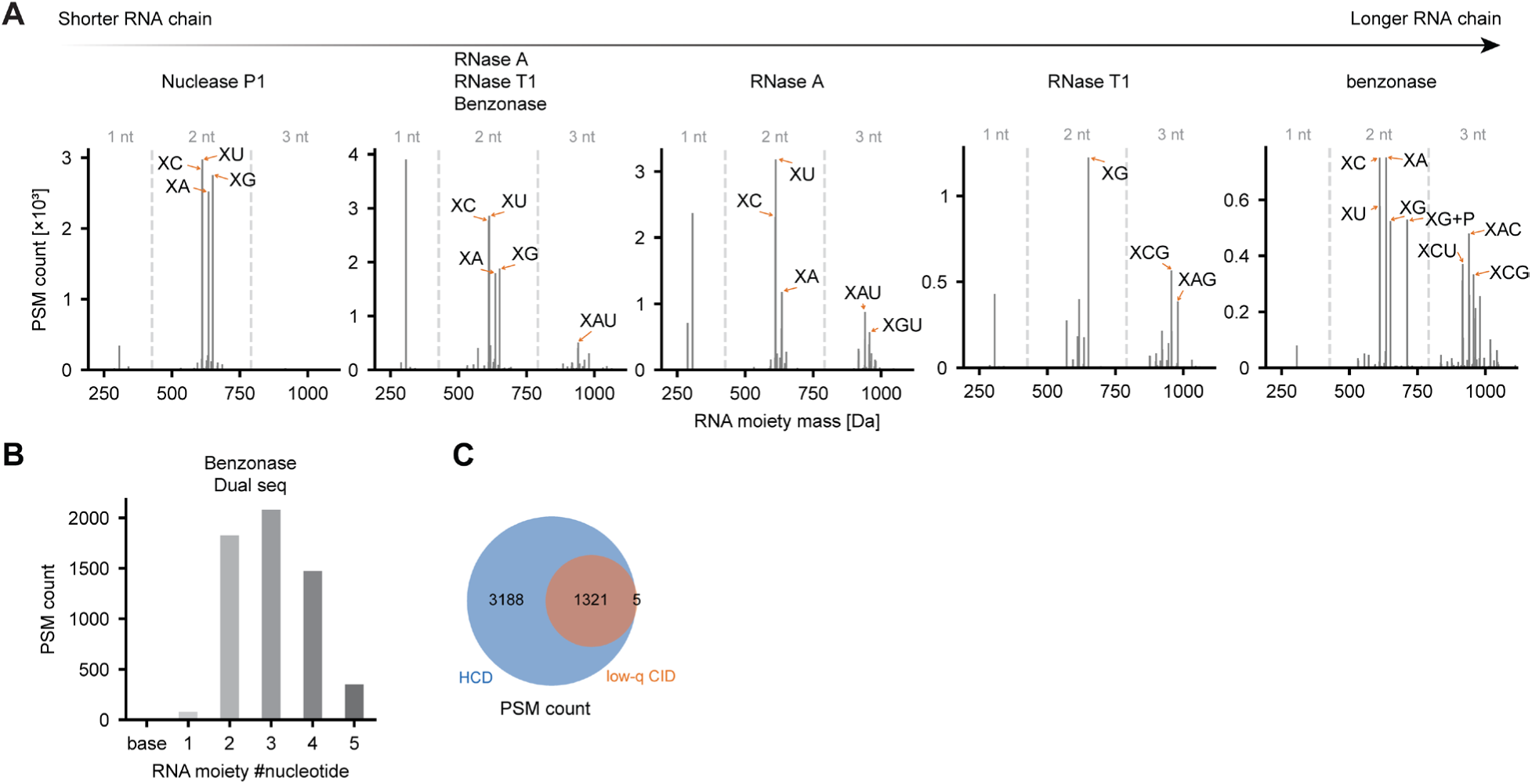
Sequencing peptide and RNA moieties of peptide-RNA crosslinks in the same MS run. **A.** Histogram of PSMs with RNA moieties identified by MSFragger offset search on samples using various nuclease combinations. Note that RNase A generated predominantly C/U-containing moieties and RNase T1 G-containing moieties, while other nonspecific nucleases produced various RNA segments. **B.** Histogram of PSMs with RNA moieties of different lengths identified by MSFragger offset search on samples using benzonase digestion, measured by the MS method shown in Figure 5B. **C.** Venn diagram comparing peptide identifications from spectra acquired with HCD and low-q CID using MSFragger offset search. Overlap indicates successfully identified peptides in both MS2 spectra of the sequential pair from the same precursor.

**Figure S6.**
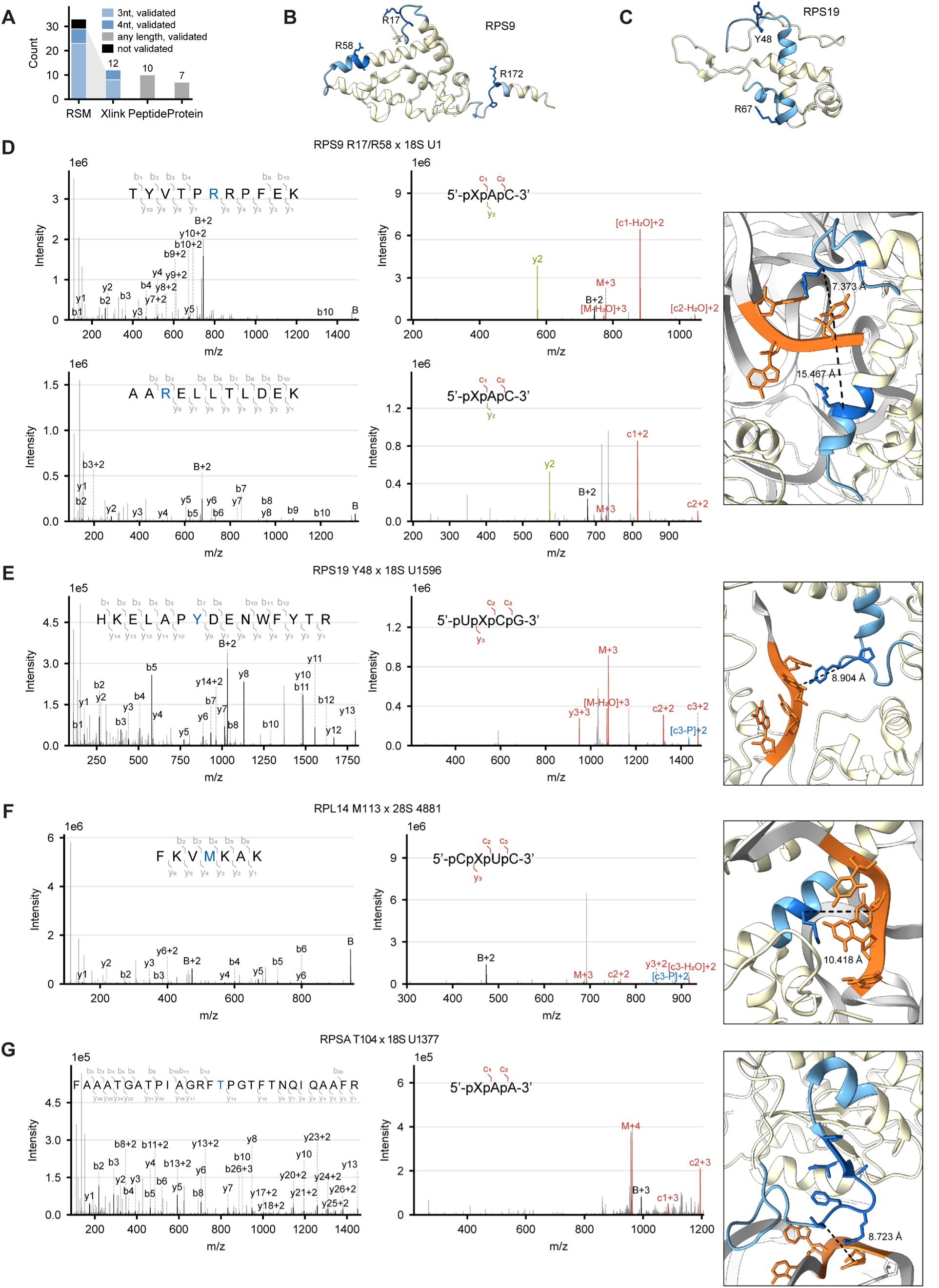
Sequenced protein-RNA crosslinks capture diverse interfaces in ribonucleoprotein complexes. **A.** Bar plot showing the numbers of RSMs, unique crosslinks, peptides and proteins validated against the human ribosome cryo-EM structure using a <20 Å distance cutoff. **B-C.** Structures of RPS9 and RPS19 in the cryo-EM structure of human 80S ribosome, with identified peptides highlighted. Light blue: identified peptide; dark blue: crosslinking sites identified by MSFragger. **D-G.** Pairs of MS2 spectra for sequential sequencing of the ribosomal crosslinks and validation in the cryo-EM structure of human 80S ribosome. Same as Figure 6A-E.

